# SHIFTR enables the unbiased and multiplexed identification of proteins bound to specific RNA regions in live cells

**DOI:** 10.1101/2023.10.09.561498

**Authors:** Jens Aydin, Alexander Gabel, Sebastian Zielinski, Sabina Ganskih, Nora Schmidt, Christina R. Hartigan, Monica Schenone, Steven A. Carr, Mathias Munschauer

**Author notes:** To whom correspondence should be addressed: Mathias Munschauer.

## Abstract

RNA-protein interactions determine the cellular fate of RNA and are central to regulating gene expression outcomes in health and disease. To date, no method exists that is able to identify proteins that interact with specific regions within endogenous RNAs in live cells. Here, we develop SHIFTR (<u>S</u>elective RNase <u>H</u>-mediated interactome framing for target RNA regions), an efficient and scalable approach to identify proteins bound to selected regions within endogenous RNAs using mass spectrometry. Compared to state-of-the-art techniques, SHIFTR is superior in accuracy, captures close to zero background interactions and requires orders of magnitude lower input material. We establish SHIFTR workflows for targeting RNA classes of different length and abundance, including short and long non-coding RNAs, as well as mRNAs and demonstrate that SHIFTR is compatible with sequentially mapping interactomes for multiple target RNAs in a single experiment. Using SHIFTR, we comprehensively identify interactions of *cis*-regulatory elements located at the 5ʹ and 3ʹ- terminal regions of the authentic SARS-CoV-2 RNA genome in infected cells and accurately recover known and novel interactions linked to the function of these viral RNA elements. SHIFTR enables the systematic mapping of region-resolved RNA interactomes for any RNA in any cell type and has the potential to revolutionize our understanding of transcriptomes and their regulation.

## INTRODUCTION

RNA carries out a plethora of functions in the cell, ranging from encoding genetic information and serving as a template for protein synthesis to dynamically regulating gene expression and controlling cellular pathways and programs. Virtually all RNA-based processes in a cell rely on interactions with specific proteins or protein complexes (1), therefore the identification of proteins that are directly bound to a target RNA can provide valuable insights into RNA function and regulation. Biochemical strategies to identify and map interactions between protein and RNA in intact cells can be divided into protein and RNA-centric approaches (2–4). Among protein-centric approaches, crosslinking and immunoprecipitation (CLIP) combined with next-generation sequencing is the most widely used technique to globally map interaction sites of a single protein of interest across the transcriptome (3). CLIP utilizes UV-crosslinking to stabilize interactions occurring in intact cells prior to cell lysis. Since UV-crosslinking creates covalent bonds between RNA and protein, denaturing purification strategies can be employed to minimize background interactions occurring in cells or after cell lysis (5–7). UV-crosslinking selectively links protein and RNA that are in direct contact, but does not efficiently crosslink indirect interactors or other biomolecules (2, 3). This is an advantage compared to more efficient chemical crosslinking strategies that have broad reactivities and covalently link DNA, protein and RNA, which results in the stabilization of indirect interactors (8). While CLIP is a powerful technique for characterizing the binding preferences of individual proteins, this approach cannot reveal the collection of proteins or protein complexes that interact with a specific endogenous RNA in intact cells. To overcome this limitation, RNA-centric interactome capture techniques were developed. Unlike protein-centric approaches that rely on the purification of individual proteins, RNA-centric methods utilize immobilized oligonucleotides that hybridize to target RNAs and enable the capture and identification of bound proteins by mass spectrometry. The first implementation of RNA interactome capture employed oligo(dT) beads to purify all polyadenylated cellular RNAs and identify proteins directly bound to mRNAs and other polyadenylated RNA species (5, 6). More recently, organic phase separation strategies have been developed to globally capture RNA-protein complexes independent of shared sequence features, such as the poly(A) tail (9–11). These methods rely on the denaturation of proteins and macromolecular complexes by guanidinium thiocyanate and phenol, followed by organic phase separation, which leads to the accumulation of RNA in the aqueous phase, while proteins accumulate in the organic phase. Unlike free RNA and free protein that separate according to their physicochemical properties during organic phase separation, covalently linked RNA-protein complexes formed as a result of UV-irradiation prior to cell lysis, cannot partition to either phase and instead accumulate in the interface (9–11). Extraction of RNA-bound proteins from UV-crosslinked interfaces after organic phase separation, followed by their identification by mass spectrometry has substantially expanded the number of known RNA-binding proteins (RBPs) (9–11).

While the aforementioned methods are powerful tools for identifying proteins bound to all cellular RNAs, they do not reveal which proteins are bound to a specific RNA. To this end, various methods that use sequence-specific oligonucleotides designed to hybridize and capture a single endogenous target RNA species have been developed (7, 12–15). Despite being technically challenging (2, 4), these approaches have been successfully used to characterize the RNA interactomes of several abundantly expressed short and long non-coding RNAs (7, 12–17), yielding important insights into their biological function. In particular, RNA antisense purification and mass spectrometry (RAP-MS) proved to be an effective tool for revealing interactomes of several different RNA types (7, 16–18). With the outbreak of the SARS-CoV-2 pandemic, RAP-MS and similar methods have also been adapted to capture and identify the proteome of host and virus that interacts with viral RNA in cells undergoing live infection (19–22).

A key limitation of virtually all methods aimed at isolating individual endogenous RNAs and their directly bound proteins is the need to generate large amounts of input material (7, 16, 19). This is due to limited crosslinking efficiency (23), incomplete RNA capture and the fact that the median mRNA copy number (∼17 per mammalian cell (24)) is orders of magnitude below the median protein copy number (∼50.000 per mammalian cell (24)), which drastically limits the amount of protein that can be purified from each cell with RNA-based capture strategies. For these reasons only highly abundant cellular RNAs have been amenable to RNA-centric interactome capture methods. Furthermore, a large number of background proteins are usually detected with RNA capture methods that target individual RNAs, impeding the selection of candidate proteins for follow-up studies. Finally, an important limitation inherent to all RNA capture approaches available to date is that proteins bound to a specific region within a larger endogenous RNA molecule cannot be identified. While capture probes can be designed to isolate a specific RNA molecule under endogenous conditions, unwanted RNA regions within this RNA molecule cannot be removed prior to protein identification without genetic engineering. Hence, the binding pattern of proteins identified with RNA capture-based strategies need to be investigated with protein-centric approaches, such as CLIP, on a one-by-one basis in follow-up studies. This is particularly limiting when *cis*-regulatory regions within an RNA molecule, such as 5ʹ and 3ʹ untranslated regions (UTRs) in mRNAs and their dynamic regulation by *trans*-acting factors are investigated. Beyond host-encoded RNAs, the genomes of RNA viruses are particularly rich in regulatory RNA elements, such as 5ʹ UTRs or leader sequences, internal ribosome entry sites (IRES), frameshifting elements, pseudoknots, or 3ʹ UTRs (25). To date, there is no method available that allows the comprehensive and unbiased mapping of region-resolved interactions between defined *cis*-acting elements in endogenous RNAs and the cellular proteome.

To overcome these limitations, we developed SHIFTR (<u>S</u>elective RNase <u>H</u>-mediated interactome framing for target RNA regions), a scalable and easy to implement low-cost approach to comprehensively identify the protein interactomes of individual endogenous RNAs or RNA elements in an unbiased fashion. SHIFTR takes advantage of recent technological breakthroughs that employ organic phase separation to globally isolate crosslinked RNA-protein complexes (9–11). Harnessing this principle, we extract UV-crosslinked RNA-protein complexes from interfaces after organic phase separation and use sequence-specific DNA probes together with RNase H to digest individual target RNAs. This results in a shift of proteins that were crosslinked to the target RNA to the organic phase. Combining SHIFTR with tandem mass tagging (TMT)-based quantitative mass spectrometry, we demonstrate the accurate and near comprehensive capture of the U1 and 7SK small nuclear ribonucleoprotein (snRNP) complexes. Comparing SHIFTR to RAP-MS for both snRNP complexes, we find that SHIFTR is superior in accurately capturing known RNP components, delivers vastly reduced background levels and requires orders of magnitude lower starting material. We establish SHIFTR workflows for targeting RNA classes of different length and abundance, including mRNAs and lncRNAs and demonstrate that SHIFTR is compatible with sequentially capturing interactomes for multiple endogenous target RNAs in a single experiment. Finally, we use SHIFTR to capture the RNA interactomes of functionally important sequence regions within the SARS-CoV-2 genome in infected human cells. We comprehensively map direct interactions between the proteome of the host cell and the authentic SARS-CoV-2 5ʹ leader as well as the viral 3ʹ UTR in cells undergoing live infection. Moreover, we complement these data with interactomes of the SARS-CoV-2 RNA genome (ORF1ab), as well as its subgenomic mRNAs (sgmRNAs) and compare the performance of SHIFTR to RAP-MS for these sequence regions. Using SHIFTR, we observe both known and novel interactions linked to the function of each targeted sequence region in the SARS-CoV-2 RNA genome. We validate interactions observed with SHIFTR at nucleotide resolution using eCLIP and find that SHIFTR uncovers binding preferences for host and viral proteins that are instructive for decoding regulatory mechanisms. Our results establish SHIFTR as a powerful new platform for region-specific RNA interactome discovery that is highly scalable, easy to implement, cost effective and yields superior near comprehensive interactome data at minimal background levels.

## RESULTS

### An organic phase separation-based strategy to define interactomes of individual RNAs

To overcome limitations of current RNA interactome capture methods, we aimed to combine the highly effective and sequence-independent isolation of RNA-protein complexes by acid guanidinium thiocyanate-phenol-chloroform (AGPC) extraction (10) with a strategy to identify proteins bound to specific RNAs or RNA regions of interest. During AGPC-based organic phase extraction of UV-crosslinked cells, covalently linked RNA-protein complexes accumulate in the interface since their physiochemical properties are incompatible with partitioning to the aqueous or organic phase (9–11). Hence, the interface accumulation of covalently linked protein-RNA complexes depends on the presence of both an RNA and a protein component. We reasoned that the selective degradation of the RNA component would release bound proteins and lead to their shift to the organic phase in a subsequent phase separation step. Proteins shifted to the organic phase upon RNA degradation can readily be extracted and identified by state-of-the-art quantitative mass spectrometry (Figure 1A). Suitable strategies to deplete RNAs or RNA regions of interest in a sequence-dependent manner include the use of specific DNA probes that hybridize to a target RNA and form RNA-DNA hybrids amenable to degradation by RNase H. We refer to this experimental strategy as SHIFTR (<u>S</u>elective RNase <u>H</u>-mediated interactome framing for target RNA regions).

**Figure 1:**
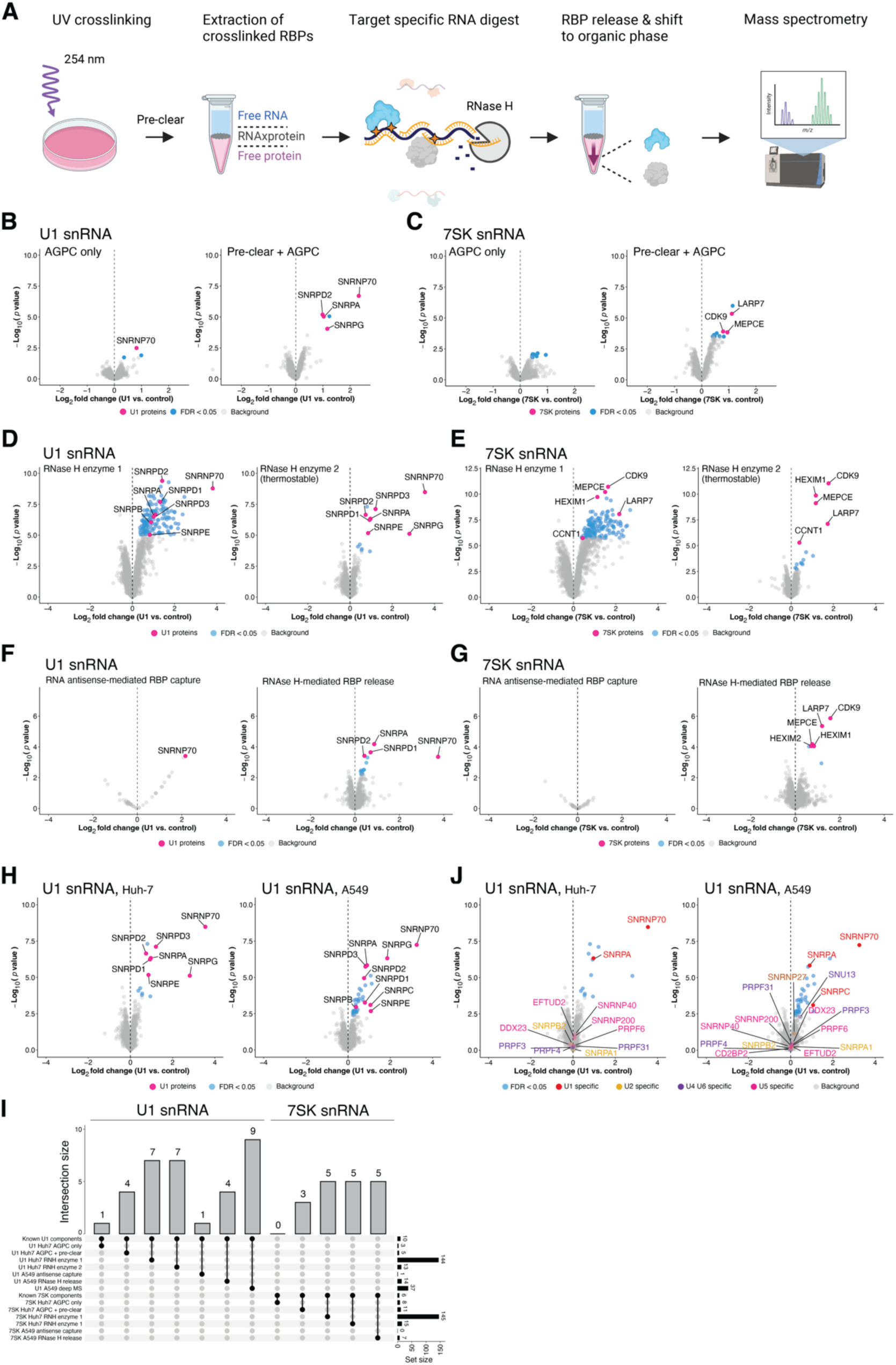
Development of a scalable, unbiased and highly efficient method to identify proteins bound to specific RNA regions in endogenously expressed RNA. **(A)** Outline of the SHIFTR workflow to identify RBPs bound to individual RNAs or RNA regions by RNase H-mediated protein release from UV-crosslinked interfaces. Illustration created with BioRender. **(B)** Quantification of proteins extracted from organic phases after targeted RNase H-mediated degradation of the U1 snRNA in UV-crosslinked interfaces by SHIFTR. Volcano plots display log_2_ fold changes comparing U1-depleted to untreated samples. Data obtained from crosslinked cells subjected to AGPC lysis without pre-treatment is shown on the left. Plot shown on the right displays data obtained from cells subjected to a pre-clearance procedure prior to AGPC treatment (Methods). Known components of the U1 snRNP complex are highlighted in pink. Experiments were performed using Huh-7 cells. RNase H (*Escherichia coli*) supplemented with BSA (enzyme 3 in Figure S1J) was used for experiments shown in **(B)** and **(C)**. **(C)** As in **(B)**, but for the 7SK snRNA. Known components of the 7SK snRNP complex are highlighted in pink. **(D)** Quantitative comparison of proteins released from UV-crosslinked interfaces when using different commercially available RNase H enzymes for SHIFTR experiments targeting the U1 snRNA. SHIFTR was performed with pre-clearing. Volcano plots display log_2_ fold changes comparing U1-depleted to untreated samples. Known components of the U1 snRNP complex are highlighted in pink. Experiments were performed using Huh-7 cells. Mass spectrometry experiments were performed with offline high pH reverse phase fractionation (Methods). **(E)** As in **(C)**, but for the 7SK snRNA. Known components of the 7SK snRNP complex are highlighted in pink. **(F)** Quantitative comparison of RNA antisense-mediated RBP capture (left) with RNase H-mediated RBP release (right) from UV-crosslinked interfaces for the U1 snRNA. Biotinylated DNA probes designed against GFP were used as a control for relative quantification. Known components of the U1 snRNP complex are highlighted in pink. Experiments were performed using A549 cells. **(G)** As in **(F)**, but for the 7SK snRNA. Known components of the 7SK snRNP complex are highlighted in pink. **(H)** Deep proteome profile of U1 SHIFTR experiments in Huh-7 (left) and A549 (right) cells using offline high pH reverse phase fractionation. Huh-7 experiment shown corresponds to data presented in **(D)** on the right. **(I)** Upset plot summarizing the enrichment of known U1 and 7SK snRNP complex components in different SHIFTR experiments with and without optimization of indicated parameters. **(J)** Analysis of SHIFTR specificity for closely related snRNP complexes. U1-specific snRNP components as well as components of other spliceosomal RNP complexes (U2, U4, U5, U6) are highlighted in different colors. Data obtained from U1 SHIFTR experiments in Huh-7 cells are shown on the left, data from A549 cells are shown on the right.

The first step to performing SHIFTR for an RNA of interest, is the design of a specific set of DNA probes for RNase H-mediated target RNA degradation. To facilitate the design of a large number of DNA probes, we developed Probe-SHIFTR (https://github.com/AlexGa/ProbeSHIFTR), a highly versatile computational tool to design, filter and select an optimized probe set for SHIFTR experiments in any species of interest. In brief, Probe-SHIFTR extracts each possible k-mer for a given target RNA and filters sequences containing stretches of polybases, repeat regions, or low complexity regions. The remaining k-mers are filtered based on sequence homology to regions in the target genome or transcriptome using BLAT (44). All filtering parameters are fully customizable. Using a greedy algorithm, Probe-SHIFTR creates multiple non-overlapping k-mer designs with minimal intra-target distances between probes. To evaluate the different probe designs, Probe-SHIFTR generates summary graphics providing information about the coverage of the target RNA region, gaps between probes, and mismatch analyses for each probe set.

### Optimization of RNase H-mediated target RNA degradation

To establish suitable conditions for the sequence-specific cleavage of target RNA-DNA hybrids by RNase H digestion, we *in vitro* transcribed the U1 RNA and synthesized a set of DNA oligonucleotides designed with Probe-SHIFTR to target different regions of U1 (Figures S1A). We used probes of 25 nucleotides in length to allow for the targeting of short sequence regions, while maintaining specificity and duplex stability. Degradation of U1 was dependent on the addition of both U1-targeting DNA probes and the RNase H enzyme (Figures S1A). When using individual probes, we observed RNA cleavage patterns consistent with the specific degradation of only the sequence regions targeted directly by the indicated DNA probes (Figure S1A). Complete degradation of the U1 RNA was achieved when using a non-overlapping probe set tiling the entire U1 RNA (Figure S1A). Since RNA extracted from UV-crosslinked cells may be less accessible for probe hybridization due to the co-purification of directly or indirectly bound proteins, we evaluated the impact of various ionic and non-ionic detergents as well as chaotropic agents that can disrupt non-covalent protein interactions on the efficiency of RNase H cleavage *in vitro* (Figures S1B, C). Detergents that did not impair RNase H activity *in vitro*, were subsequently tested with interfaces extracted from UV-crosslinked Huh-7 cells after organic phase separation. We observed improved depletion of the U1 RNA when adding a combination of 3 non-ionic detergents, while ionic detergents and chaotropic agents reduced the efficiency of U1 depletion, as estimated by RT-qPCR (Figure S1D). Hence, we selected the non-ionic detergent-supplemented buffer system to help improve the solubility and accessibility of extracted cellular RNA-protein complexes and achieve optimal protein release. To further maximize the efficiency of target RNA digestion, we used different concentrations of DNA probes and evaluated the efficiency of U1 depletion (Figure S1E). We observed the highest U1 depletion with a concentration of 5 μM, with no considerable unspecific degradation of other RNA targets, such as 7SK.

### Pre-treatment of UV-crosslinked cells enhances RNA interactome capture by SHIFTR

Next, we optimized the purification of UV-crosslinked RNA-protein complexes from living cells by AGPC extraction. While the clean purification of interfaces is a crucial step in this workflow, prior work already indicated that the co-purification of DNA-binding proteins (9), as well as membrane components (11) and glycosylated proteins (10) by organic phase extraction can result in the enrichment of irrelevant proteins. Additionally, the high complexity of UV-crosslinked cell pellets directly subjected to organic phase extraction may interfere with the clean partitioning of RNA-protein complexes in interfaces and hinder efficient RNA digestion as well as protein release. We speculated that an additional clearance and purification procedure may enhance the sequence-specific release of proteins bound by target RNA sequences from interfaces after organic phase extraction. We subjected cells to detergent-based lysis followed by a DNase treatment to remove contaminating DNA and a centrifugation step to remove membranes and cell debris (Methods). We selected the well-characterized small nuclear RNAs (snRNAs) U1 and 7SK for proof-of-principle experiments and compared the effective capture of known protein components with and without subjecting UV-crosslinked cells to the described pre-clearance and DNase treatment. We first confirmed the specific depletion of target RNAs after interface extraction and RNase H cleavage by RT-qPCR (Figure S1F). Next, we performed western blot analysis and observed specific enrichment of known components of the U1 and 7SK complexes in the respective SHIFTR experiments (Figure S1G). To minimize the isolation of background proteins, we extracted UV-crosslinked interfaces and performed up to four consecutive organic phase separations while monitoring the isolation of background as well as U1 target proteins. As shown in Figure S1H, little free background protein was detected in the organic phase after 3-4 phase separations. At the same time the specific release of the U1 interacting protein SNRNP70 was not negatively impacted by multiple successive organic phase extractions (Figure S1I). To confirm the effective solubilization of intact total RNA and assess whether the pre-clearance and DNase step has any impact on RNA quality, we released crosslinked RNA from interfaces by proteinase K treatment and analyzed RNA integrity by capillary electrophoresis. We noted that in the absence of UV crosslinking interfaces from pre-cleared cell pellets contained less contaminating RNA. In UV-crosslinked samples, the 28S rRNA peak appeared slightly more degraded when pre-clearing was performed (Figure S1K). In contrast, the 18S rRNA peak did not show any sign of enhanced degradation with pre-clearing (Figure S1K).

To quantitatively compare the specific enrichment of known U1 and 7SK components in SHIFTR experiments with or without the described pre-clearance and DNase treatment, we performed quantitative TMT-based mass spectrometry analyses. We used 10 million UV-crosslinked Huh-7 cells for each SHIFTR experiment. As shown in Figures 1B-C, the combined pre-clearing and DNase treatment substantially improved the recovery of known U1 and 7SK proteins with 4 components of the U1 snRNP complex (SNRNP70, SNRPG, SNRPA, SNRPD2) and 3 known 7SK components (LARP7, MEPCE, CDK9) displaying a statistically significant enrichment. Beyond the known components of the U1 snRNP, we found only one other protein among significantly enriched candidates in U1 SHIFTR experiments (Table S1). Importantly, this protein (SF3A1) was previously shown to directly bind the stem-loop 4 (SL4) of U1 (45) and likely represents a true U1 complex member. In the absence of pre-clearing and DNase treatment, we found only one known U1 component (SNRNP70) and no known 7SK components that met our statistical significance cutoff (Figures 1B-C, Table S1). Hence, the pre-clearance and DNase treatment substantially improves the recovery of known RNA-protein complexes and appears particularly effective for RNAs less abundant than U1, including the 7SK snRNA, for which no known interactors were found in the absence of pre-clearing.

### Thermostable RNase H displays superior specificity and effective on-target protein release in SHIFTR experiments

Next, we evaluated the specificity and efficacy of different commercially available RNase H enzymes for releasing RNA-bound proteins in SHIFTR experiments. We focused on RNase H preparations that do not contain bovine serum albumin (BSA) or other protein additives (Figure S1J) and compared thermostable RNase H with standard RNase H from two different vendors. Using standard RNase H, we observed an unexpectedly large number of enriched proteins when targeting the U1 or 7SK snRNAs with specific DNA probes (Figures 1D-E, S1L-M). The majority of these proteins were RBPs unrelated to the U1 or 7SK snRNP (Table S1), suggesting unspecific and probe-independent RNA degradation with this enzyme preparation. In contrast, thermostable RNase displayed high specificity and selectivity for the targeted RNA-protein complexes, as indicated by the effective release of known U1 and 7SK components (Figures 1D-E, S1L-M). We next performed RNA-sequencing to compare control samples to U1 SHIFTR samples treated with thermostable RNase H and observed multiple U1 gene copies as the only differentially expressed genes (Figure S1N). Hence, we used thermostable RNase H for all subsequent experiments. We also tested whether treating RNase H-digested samples with the exoribonuclease Xrn1 to remove residual RNA fragments crosslinked to the released proteins would enhance the shift of known U1 or 7SK complex members to the organic phase, but observed no detectable change by western blot analysis (Figure S1O).

### RNase H-mediated protein release from crosslinked interfaces is superior to RNA antisense purification

We next evaluated if the degradation of target RNA sequences by RNase H, which leads to a shift of the released proteins to the organic phase, is more effective than a direct capture of target RNAs from interface extracts using biotinylated DNA antisense probes. To this end, we extracted interfaces from 10 million UV-crosslinked cells and either followed the SHIFTR workflow as described above, or solubilized interfaces in RAP-MS hybridization buffer (7). We added pools of biotinylated antisense probes designed to hybridize to the U1 or 7SK snRNAs to the fully solubilized interfaces and followed the previously described RAP-MS protocol for antisense capture (7). We monitored the effective enrichment or depletion of target RNAs by RT-qPCR (Figure S1P) and quantitatively analyzed the U1 and 7SK RNA interactomes obtained from interfaces subjected to RAP-MS or SHIFTR workflows (Figures 1F-G). While we were able to readily identify known U1 and 7SK complex members using the SHIFTR approach, RNA antisense-mediated purification of U1 and 7SK bound proteins from UV-crosslinked interfaces yielded only a single known complex member (SNRNP70) in the case of U1 (Figures 1F). For the 7SK snRNA, the RNA antisense purification approach did not yield any known complex members among significantly enriched proteins in quantitative mass spectrometry experiments (Figure 1G). Hence, with the scale and experimental setup used, SHIFTR is superior to the combined organic phase separation and RNA antisense purification strategy described above.

### An optimized MS strategy for comprehensive recovery of U1 and 7SK RNA interactomes

Following the detailed optimization of the SHIFTR workflow, we set out to improve the comprehensive identification of RNA-bound proteins released from UV-crosslinked interfaces. To this end, we included an offline high pH reverse phase fractionation step prior to sample injection into the mass spectrometer to reduce sample complexity and improve the resolution of obtained mass spectra. As expected, this increased the number of identified proteins from 1642 proteins across all samples without fractionation to 3695 proteins with offline fractionation (Tables S1-S2). With this modification, we recovered one additional U1 snRNP core protein and two additional 7SK binding proteins in the same SHIFTR samples described above. We also noted an increase in the number of peptides detected for each identified protein, thus increasing confidence in our measurements. In total, we identified 7 out of 10 core interactors of the U1 snRNP (Figures 1H (left), I) and 5 out of 6 core interactors of the 7SK snRNP in Huh-7 cells (Figures 1I, S1Q (left)). Our experimental conditions were readily compatible with an alternative cell model (here, A549 cells) where we identified 9 out of 10 known interactors of the U1 snRNA (Figures 1H (right), I) and 5 out of 6 core interactors of the 7SK snRNA (Figures 1I, S1Q (right)) without any modifications to the protocol (Tables S1-S2). Remarkably, SHIFTR is not only compatible with the near comprehensive capture of known snRNP components across different cell types, but also displays superb specificity for the targeted snRNP, as indicated by the selective enrichment of U1-specific snRNP components, while proteins specific for the closely related U2, U4, U5 and U6 complexes, were not enriched in either cell type (Figure 1J).

### SHIFTR is superior to RAP-MS in a systematic side-by-side comparison

Having established an optimized SHIFTR protocol, we systematically compared the performance of SHIFTR to RAP-MS as the state-of-the-art method for RNA interactome capture for individual cellular RNAs (4, 16, 19). We performed RAP-MS for the U1 and 7SK snRNP complexes (Figures S2A-B) using 400 million UV-crosslinked cells, which is in line with previous reports (7, 16, 17, 19). A direct comparison of significantly enriched proteins in SHIFTR and RAP-MS experiments revealed that both methods successfully identified a comparable number of known U1 (Figures 2A-B, Table S3) and 7SK components (Figures 2C-D, Table S3). Strikingly however, many proteins not directly linked to known 7SK functions were significantly enriched in RAP-MS experiments, while SHIFTR experiments were virtually free of background interactions (Figures 2A-D, Table S3). Leveraging our detailed knowledge about the composition of both the U1 (46) and 7SK (47) snRNPs, we evaluated the accuracy of both methods for capturing the respective RNP complexes using the F1-score, which combines precision and recall in a single metric. SHIFTR yielded much higher F1-scores for both RNP complexes (Figure 2E). Similarly, SHIFTR consistently outperforms RAP-MS for both the U1 and 7SK complexes at all significance levels when comparing precision and recall statistics (Figures 2F-G, S2C). In line with these results, we also found that the fraction of true positive U1 and 7SK complex members among significantly enriched proteins was much higher for SHIFTR (Figure S2D). Together, our experiments show that SHIFTR yields superior signal to noise ratios compared to RAP-MS and delivers more accurate RNA interactomes from a fraction of the input material, which dramatically expands the scalability and applicability of RNA interactome capture methods.

**Figure 2:**
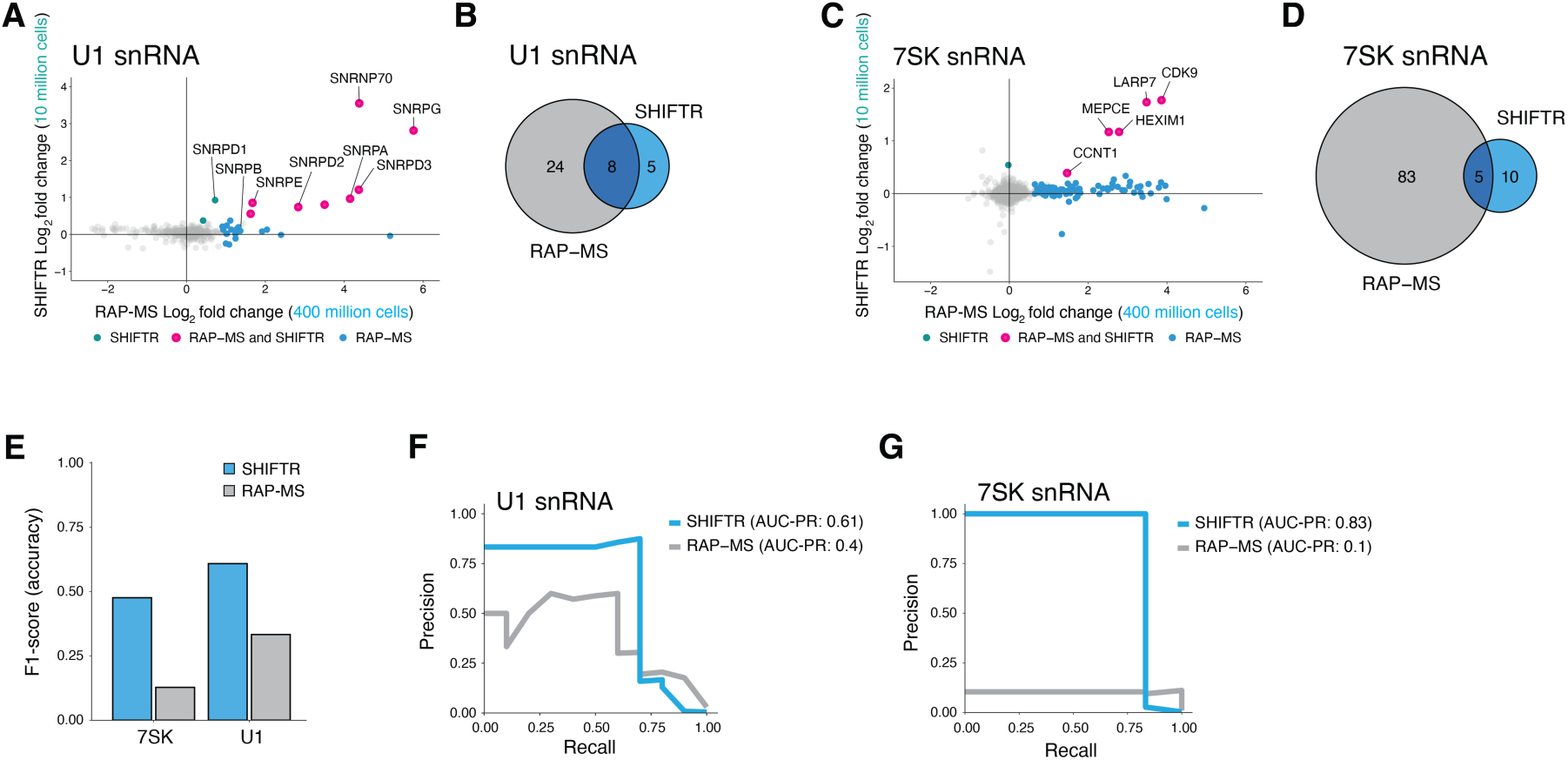
Side-by-side comparison of the performance of SHIFTR and RAP-MS for the U1 and 7SK snRNPs. **(A)** Fold change correlation plot comparing the enrichment of U1-interacting proteins identified with SHIFTR (y-axis) and RAP-MS (x-axis) methods. Proteins significantly enriched with both methods are highlighted in pink, background proteins are shown in grey. Proteins only detected with SHIFTR are shown in teal and proteins only detected with RAP-MS are shown in blue. **(B)** Venn diagram comparing proteins significantly enriched in RAP-MS and SHIFTR experiments targeting the U1 snRNA. **(C)** As in **(A)**, but for the 7SK snRNA. **(D)** As in **(B)**, but for the 7SK snRNA. **(E)** Comparison of the F1-score obtained for the U1 and 7SK snRNAs when using either SHIFTR (blue) or RAP-MS (grey) to identify directly bound proteins. **(F)** PR curve displaying precision and recall values for SHIFTR and RAP-MS experiments when targeting the U1 snRNA. Area under the PR curve (AUC-PR) is analyzed for SHIFTR (0.61) and RAP-MS (0.40). **(G)** As in (F), but for the 7SK snRNA. AUC-PR is analyzed for SHIFTR (0.83) and RAP-MS (0.10).

### Sequential release of multiple target RNA interactomes with a streamlined SHIFTR workflow

To unlock the full potential of SHIFTR, we proceeded with targeting coding and non-coding cellular RNAs that differ in length and abundance from the previously interrogated snRNAs. We initially used the same amount of input material that was sufficient to comprehensively capture the U1 and 7SK snRNP complexes and targeted the nuclear lncRNA MALAT1, as well as the actin beta (ACTB) mRNA. These initial experiments yielded few enriched proteins (data not shown), suggesting that the amount of input material was likely insufficient for the detection of proteins crosslinked to these less abundant RNAs by mass spectrometry. Since scaling up the SHIFTR workflow requires increasing the volumes of organic solvents beyond the scale of what can be readily handled using standard microfuge tubes and benchtop instruments, we decided to combine SHIFTR with a widely used poly(A)-based RNA interactome capture procedure (5, 6, 48, 49) (Figure 3A). This approach concentrates crosslinked RNAs in a small volume and avoids introducing technical variation by extensive partitioning of samples into multiple microfuge tubes, which would also be incompatible with processing many samples in parallel, thus limiting scalability. We used 150 million UV-crosslinked A549 cells and replaced the pre-clearance and DNase treatment described above with a simplified poly(A)-based RNA interactome capture step (Methods). This modification has multiple important advantages. First, as demonstrated above, a pre-clearance or sample purification step prior to organic phase extraction enhances the specific release of RNA bound proteins in SHIFTR experiments (Figures 1B-C). Second, reaction volumes are reduced such that a single microfuge tube is sufficient for subsequent manipulation steps, ensuring scalability of the approach. Third, with an initial poly(A)-based RNA interactome capture step, the scale of SHIFTR experiments can be adjusted such that multiple sequential RNase H digestions can be performed using the same interface extract to release the proteins bound to different RNA targets in a sequential fashion (Figure 3A). To demonstrate this, we performed organic phase extraction after poly(A)- based RNA interactome capture and digested the ACTB mRNA using sequence-specific DNA probes together with RNase H. As expected, we observed specific depletion of ACTB only when sequence-specific DNA probes were added (Figure S3A). After completing the SHIFTR workflow for ACTB, we used the same interface after organic phase separation to digest the nuclear lncRNA MALAT1 with sequence-specific DNA probes and RNase H, which resulted in a strong reduction of MALAT1 RNA levels only when specific DNA probes were added (Figure S3B). We completed the SHIFTR workflow for these two cellular RNAs (Figure S3C) and subjected resulting samples to quantitative TMT-based LC-MS/MS analysis. Since the identification of direct RNA-protein interactomes is particularly challenging for individual mRNAs due to their limited cellular copy number, we conducted an additional proof-of-principle experiment targeting the GAPDH mRNA using SHIFTR. In each case, we observed many annotated RBPs among significantly enriched proteins (Figures 3B-D, Table S4), validating our approach. To our knowledge, these data represent the first global RNA-protein interactomes of individual mRNAs collected in unmodified cells under endogenous conditions. To further benchmark the performance of SHIFTR, we generated an additional MALAT1 RNA interactome using RAP-MS as a reference (Table S3, Figure S3D). Comparing MALAT1 SHIFTR and RAP-MS data (Figures 3E-F), we found 11 overlapping proteins, including HNRNPC, a functionally supported MALAT1 interacting protein (50), which was among the most significantly enriched proteins using both methods. Assuming that proteins independently identified with both methods likely represent true direct interactors, our complementary experimental strategy defines a core high-confidence RNA-protein interactome of MALAT1 that may be harnessed for future functional or mechanistic experiments.

**Figure 3:**
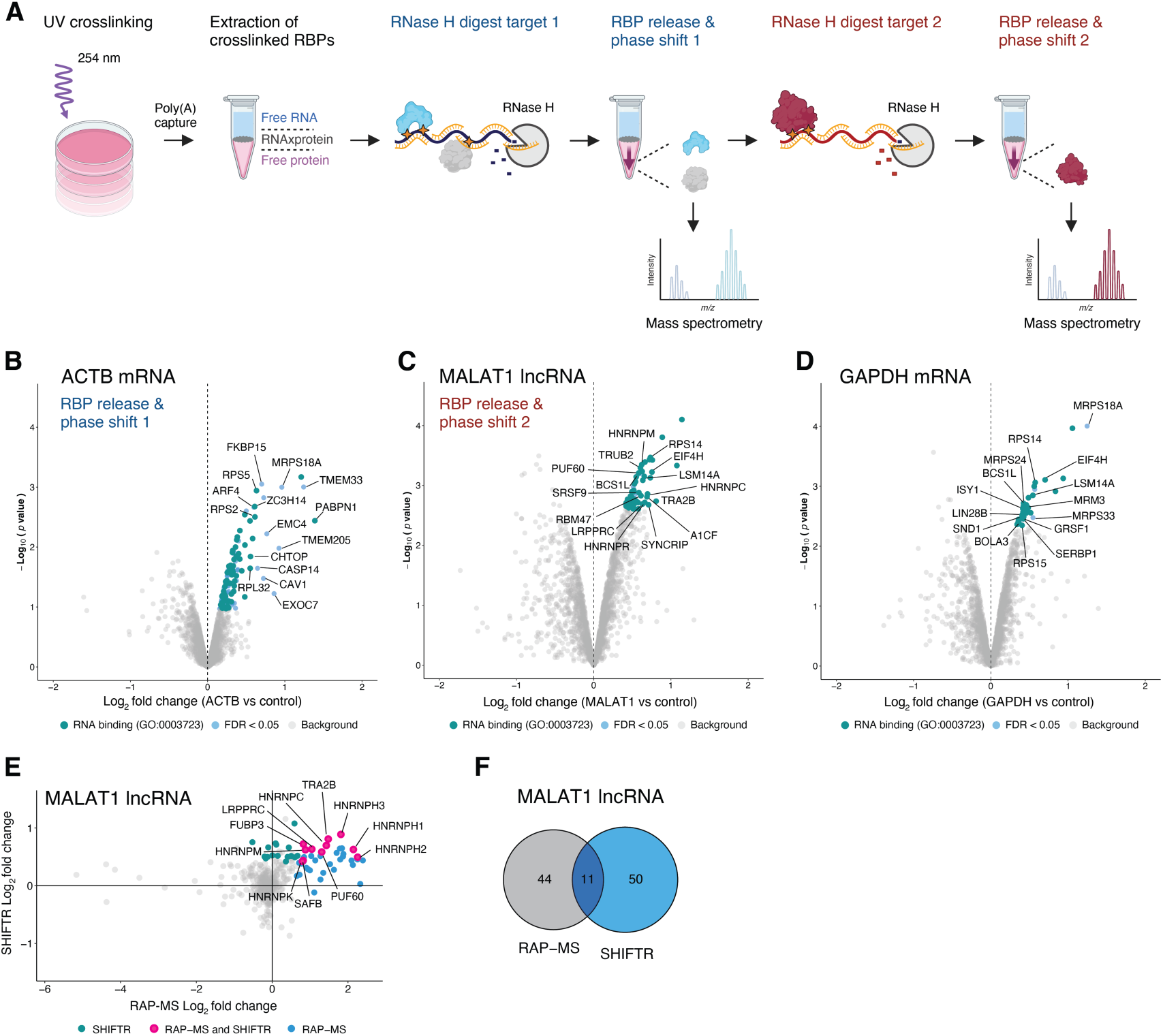
Sequential targeting of different RNA types with a streamlined SHIFTR workflow. **(A)** Outline of a streamlined SHIFTR workflow incorporating poly(A)-based bulk RNA interactome capture for pre-clearing, as well as multiple RNase H digestion steps for the sequential release of RBPs bound to specific individual RNAs in a multiplexed fashion. Illustration created with BioRender. **(B)** Quantification of proteins extracted from the organic phase after targeted RNase H-mediated degradation of the ACTB mRNA in UV-crosslinked interfaces by SHIFTR. Volcano plot displays log_2_ fold changes comparing ACTB-depleted to untreated samples. Significantly enriched proteins are displayed in blue and significantly enriched proteins annotated as RNA-binding protein (GO:0003723) are highlighted in teal. **(C)** As in **(B)**, but for the MALAT1 lncRNA. Note that the same interface used for depleting the ACTB mRNA, was subsequently used for the depletion of MALAT1 and the extraction of MALAT1-bound RBPs. **(D)** As in **(B)**, but for the GAPDH mRNA and using a separate UV-crosslinked interface. **(E)** Fold change correlation plot comparing the enrichment of MALAT1-interacting proteins identified with SHIFTR (y-axis) and RAP-MS (x-axis) methods. Proteins significantly enriched with both methods are highlighted in pink, background proteins are shown in grey. Proteins only detected with SHIFTR are shown in teal and proteins only detected with RAP-MS are shown in blue. **(F)** Venn diagram comparing proteins significantly enriched in RAP-MS and SHIFTR experiments targeting the MALAT1 lncRNA.

### SHIFTR defines the interactomes of genomic and subgenomic SARS-CoV-2 RNAs in infected cells

Due to their limited size, the genomes of RNA viruses are particularly rich in *cis*-regulatory RNA elements that recruit *trans*-acting factors of virus and host to facilitate RNA replication and virus propagation. Despite the functional importance of these *cis*-acting RNA elements, the identification of proteins that bind to specific sequence regions in authentic viral RNAs has not been possible with available technologies. Hence, we selected SARS-CoV-2 as a model virus to demonstrate the power of SHIFTR for revealing sequence region-resolved RNA-protein interactions. SARS-CoV-2 is an enveloped, positive-sense, single-stranded RNA virus. Upon infection of a host cell, the virus deploys its 5ʹ-capped and 3ʹ-polyadenylated RNA genome to express a set of viral proteins crucial for replication and virus propagation using the protein synthesis machinery of the host. During infection, SARS-CoV-2 also generates a nested set of 5ʹ and 3ʹ co-terminal subgenomic mRNAs that encode structural and accessory viral proteins. Similar to other RNA viruses, SARS-CoV-2 contains non-coding regulatory RNA elements at the 5ʹ and 3ʹ-end of all viral RNAs that are important for viral replication (51). To identify proteins that directly bind specific regions of the SARS-CoV-2 genome, including the regulatory RNA elements at the 5ʹ and 3ʹ-ends, we infected A549^ACE2^ cells (52) with SARS-CoV-2 and performed SHIFTR experiments at 24 hours post infection (hpi). We first targeted the ORF1ab region in SARS-CoV-2 RNA, which is uniquely present in viral RNA genomes, but not in sgmRNAs. In separate experiments we designed a probe set overlapping all sgmRNAs (S, ORF3a, E, M, ORF6, ORF7a, ORF7b, ORF8, N, ORF10). As shown in Figure S4A, we observed a strong depletion of the targeted viral RNA regions. Of note, the ORF1ab targeting probe set contained probes designed to deplete the 5ʹ leader sequence, which was readily observed in our experiment (Figure S4A). Similarly, probe sets used for depleting subgenomic mRNAs contained probes that target both the 5ʹ leader and the viral 3ʹ UTR, which are both present in all sgmRNAs (Figure S4A). We next proceeded with analyzing the proteins released from crosslinked interfaces by ORF1ab and sgmRNA depletion using quantitative TMT-based LC-MS/MS.

Among the 48 significantly enriched proteins in ORF1ab SHIFTR experiments, ∼67 % were annotated as RBPs based on GO annotations (Figure 4A, Table S5). Since the ORF1ab region is only present in the RNA genomes of SARS-CoV-2, we compared ORF1ab SHIFTR data to RAP-MS data that we generated for the same sequence region (Figures 4B, S4B) (53). For RAP-MS we used 160 million SARS-CoV-2 infected and UV-crosslinked A549^ACE2^ cells (24 hpi), while SHIFTR was performed with 10 million cells. Among significantly enriched factors, 15 proteins were identified with both methods, representing high-confidence ORF1ab RNA interactors. Focusing on specific candidates, we noticed the nucleoprotein (N) of SARS-CoV-2 (Figure 4A-B, Table S5). Compared to other viral RNA regions analyzed in this work, N is most strongly enriched when targeting the ORF1ab region (Table S5). Importantly, N is known to densely cover the viral RNA genome and is required for genome packaging (54). Hence, the binding pattern observed in SHIFTR experiments is consistent with the known function of N in the viral replication cycle. Similarly, we found the host antiviral protein ZAP (ZC3HAV1) strongly enriched among ORF1ab-bound proteins (Figure 4B, Table S5). Prior work showed that ZAP interacts with the frameshift element (FSE) in ORF1ab *in vitro* and regulates the SARS-CoV-2 –1 frameshifting efficiency (55), which is in line with the enrichment of ZAP in ORF1ab SHIFTR experiments observed here.

**Figure 4:**
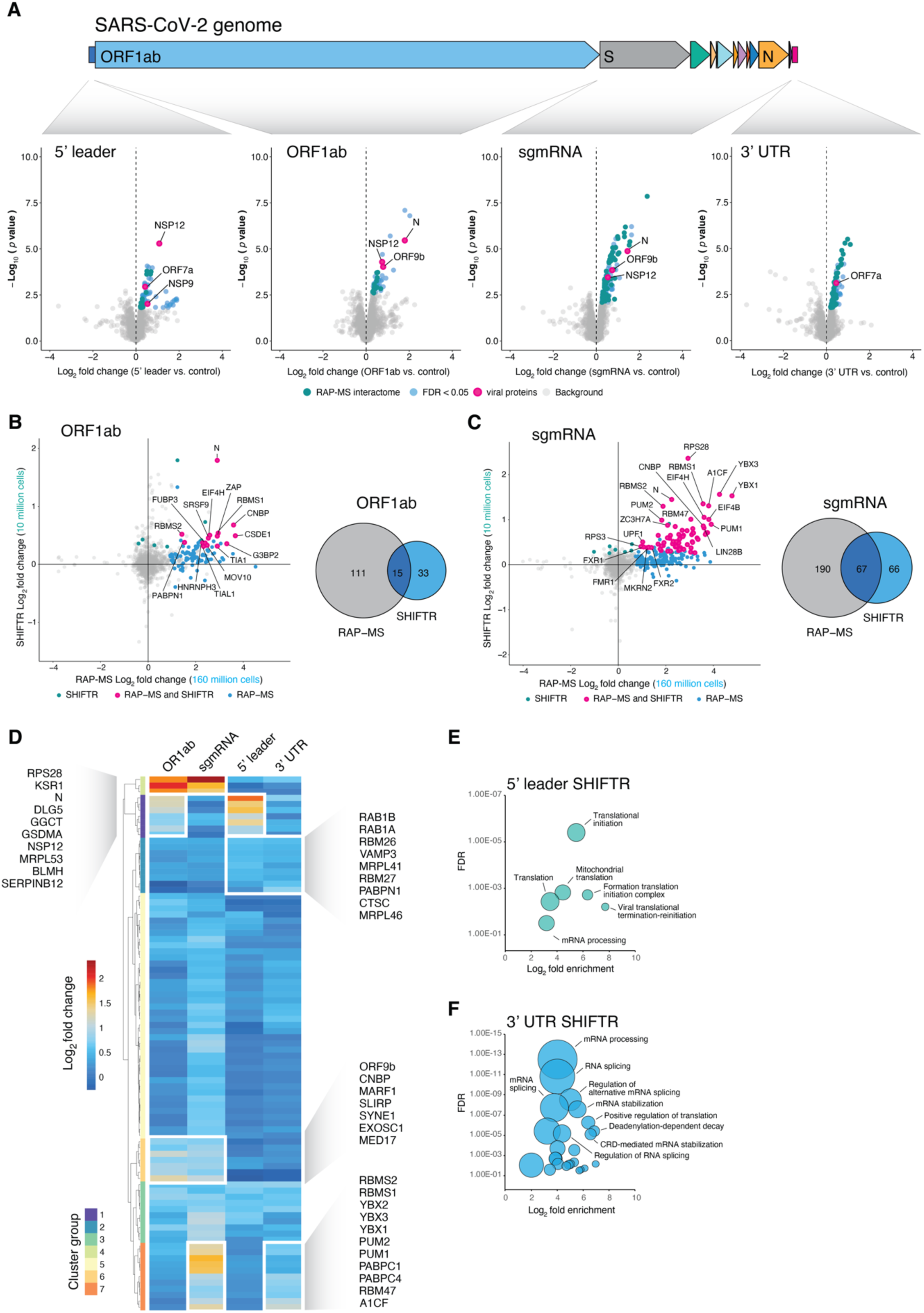
SHIFTR delivers a high-resolution view on the protein interactions of different sequence regions in the authentic SARS-CoV-2 RNA genome. **(A)** Schematic illustration of the different sequence regions in the SARS-CoV-2 genome interrogated with SHIFTR. Zoom in views display the proteins identified with SHIFTR to bind to the respective RNA region. Volcano plot displays log_2_ fold changes comparing interfaces digested for the indicated SARS-CoV-2 RNA region (from left to right: 5ʹ leader, ORF1ab, sgmRNA, 3ʹ UTR) relative to untreated interfaces. Significantly enriched proteins are displayed in blue and significantly enriched proteins also discovered in RAP-MS-based SARS-CoV-2 RNA interactome data (19) are highlighted in teal. Viral proteins are highlighted in pink. **(B)** Fold change correlation plot comparing the enrichment of proteins identified as interactors of the ORF1ab region of SARS-CoV-2 with SHIFTR (y-axis) and RAP-MS (x-axis) methods. Proteins significantly enriched with both methods are highlighted in pink, background proteins are shown in grey. Proteins only detected with SHIFTR are shown in teal and proteins only detected with RAP-MS are shown in blue. Venn diagram comparing proteins significantly enriched in RAP-MS and SHIFTR experiments targeting the ORF1ab region of SARS-CoV-2 is shown on the right. RAP-MS data is part of separate study (53). **(C)** As in **(B)**, but for the S to ORF10 region, encoding SARS-CoV-2 sgmRNAs. **(D)** Heatmap displaying the enrichment of candidate proteins when targeting the indicated SARS-CoV-2 RNA regions with SHIFTR. Only candidates significantly enriched in at least one SHIFTR experiment and with a log_2_ fold change > 0.5 are displayed. White boxes highlight selected clusters of interest. **(E)** GO enrichment analysis of proteins significantly enriched in SARS-CoV-2 5ʹ leader SHIFTR experiments. Circle sizes scale to the number of detected proteins. **(F)** As in **(E)**, but for proteins significantly enriched in SARS-CoV-2 3ʹ UTR SHIFTR experiments.

We next analyzed proteins identified in sgmRNA SHIFTR experiments and found that ∼80 % of the 133 significantly enriched proteins were annotated as RBPs (Figure 4A, Table S5). When comparing interactors of sgmRNAs identified using SHIFTR or RAP-MS, we observed 67 shared high-confidence interactors, representing >50 % of all significantly enriched proteins identified with SHIFTR (Figures 4C, Table S5). These results are in line with our previous observations that SHIFTR yields comparable RNA interactome data from a fraction of the input material needed for RAP-MS. Consistent with our RAP-MS data (19), we found several proteins of the La-related protein (LARP) family, including LARP1 and LARP4 among the strongest sgmRNA interactors (Table S5). We previously demonstrated that LARP1 plays an important role in restricting SARS-CoV-2 RNA replication, likely by repressing the translation of viral mRNAs (19). Beyond the LARP family, we also observed a strong enrichment of CNBP in sgmRNA SHIFTR experiments (Table S5), which is consistent with earlier reports suggesting that CNBP directly binds SARS-CoV-2 RNA and restricts SARS-CoV-2 replication in human cells (19, 56). Together, the interaction partners and binding preferences uncovered with SHIFTR are consistent with the known function and regulation of the targeted viral sequence regions.

### SHIFTR uncovers interactions of the 5ʹ and 3ʹ terminal regions of the authentic SARS-CoV-2 RNA genome in infected cells

We next focused on uncovering interactions of short *cis*-regulatory RNA elements located at the 5ʹ and 3ʹ-ends of the authentic SARS-CoV-2 RNA genome. While RAP-MS can be readily modified to biochemically separate RNA molecules containing different sequence regions, such as genomic and subgenomic SARS-CoV-2 RNAs, it is not possible to interrogate individual sequence elements within endogenous RNAs using antisense capture-based approaches. Hence, with technologies available to date, it has not been possible to globally characterize the interactions of the 5ʹ and 3ʹ-terminal regions of authentic viral RNA genomes in infected cells, despite their critical importance for viral replication and host immune sensing (57). We performed SHIFTR in SARS-CoV-2 infected cells as described above and targeted the 5ʹ leader sequence as well as the viral 3ʹ UTR in separate experiments. In each case we observed specific depletion of the targeted RNA region, while other regions in the viral genome remained unperturbed (Figure S4A). We subjected the proteins released by SHIFTR to quantitative TMT-based LC-MS/MS analysis as described above.

A global analysis of the proteins bound to the four different SARS-CoV-2 sequence regions targeted in this work, revealed similarities as well as notable differences in their respective RNA interaction profiles (Figure 4D, Table S5). The protein interaction profile of the SARS-CoV-2 3ʹ UTR for instance, appears overall well represented in the RNA interactome of sgmRNAs (Figure 4D). In contrast, translational regulators were mostly found within sgmRNAs and the ORF1ab sequence (Table S5), while 3ʹ UTRs were more strongly bound by RNA processing factors and proteins known to regulate RNA stability (Table S5). We next compared the compendium of human proteins identified in SHIFTR experiments targeting different SARS-CoV-2 RNA regions to SARS-CoV-2 host factors identified based on functional genetic screens and interactome studies in various different cell types (58). Of the 186 unique human proteins collectively identified in SHIFTR experiments targeting different SARS-CoV-2 RNA regions, 92 were previously identified as SARS-CoV-2 RNA binders (Figure S4D). SHIFTR also uncovered 94 proteins that were not previously recognized to bind SARS-CoV-2 RNA and 8 of these factors are functionally relevant for SARS-CoV-2 infection based on genetic evidence (58) (Figure S4D). Hence, in addition to revealing binding preferences of known SARS-CoV-2 RNA binders, SHIFTR uncovers novel interactions, including interactions that play functionally important roles in SARS-CoV-2 infection.

### Insights into interactions and functions of the SARS-CoV-2 5ʹ leader

Focusing on specific interactions of the 5ʹ leader, we found 60 significantly enriched proteins in SHIFTR experiments, of which ∼50 % are annotated as RBPs (Figure 4A, Table S5). Among significantly enriched proteins, we observed several translation initiation factors (EIF4B, EIF4H, EIF3G, EIF3CL, EIF3D, EIF3A), which is consistent with the known role of the 5ʹ leader in recruiting machinery of the host cell to initiate viral mRNA translation (51, 59) (Table S5). A GO enrichment analysis confirmed the overrepresentation of proteins linked to translational initiation among 5ʹ leader interactors (Figure 4E). Beyond these host proteins, we observed several viral proteins including the RNA-dependent RNA polymerase NSP12, as well as the viral RNA-binding protein NSP9 among significantly enriched 5ʹ leader interactors (Figure 4A). These interactions are consistent with key roles of the 5ʹ leader in viral RNA biogenesis. Importantly, recent *in vitro* reconstitution experiments demonstrated that NSP9 can be covalently linked to the 5ʹ leader of SARS-CoV-2 (60, 61). Intriguingly, the covalent linkage of NSP9 to the 5ʹ end of SARS-CoV-2 RNA suggests a role in priming of viral RNA synthesis and/or a function in the capping mechanism of SARS-CoV-2 RNAs (60–63). The detection of NSP9 as a significantly enriched viral RNA binder in SHIFTR experiments targeting the 5ʹ leader is highly consistent with these observations and the emerging functions of NSP9 in the biogenesis of viral RNA.

### Insights into interactions and functions of the SARS-CoV-2 3ʹ UTR

Finally, when analyzing SHIFTR experiments targeting the SARS-CoV-2 3ʹ UTR, we found 99 significantly enriched proteins, of which ∼75 % are annotated as RBPs (Figure 4A, Table S5). We observed an overrepresentation of GO terms linked to the regulation of mRNA processing and stability through binding of 3ʹ UTR elements (Figure 4F, Table S5). We noted many well-known post-transcriptional regulators, such as the PUM1 and PUM2 proteins (Figure 4D, Table S5) that preferentially bind and regulate 3ʹ UTR sequences in host mRNAs (64). Similarly, several proteins that regulate mRNA stability by binding near the poly(A) tail of mRNAs were among the most strongly enriched candidates in the SARS-CoV-2 3ʹ UTR (Figure 4D, Table S5). These include the cytoplasmic poly(A)-binding proteins PABPC1 and PABPC4, the poly(ADP-ribose) polymerase PARP12, and PATL1, a regulator of mRNA deadenylation-dependent decapping (Figure 4D, Table S5). Preferential binding of these host-encoded post-transcriptional regulators to the 3ʹ terminal region of SARS-CoV-2 is consistent with known roles of the viral 3ʹ UTR in controlling RNA stability and translation (51). Among viral proteins significantly enriched in SHIFTR experiments targeting the SARS-CoV-2 3ʹ UTR, we found ORF7a, a viral transmembrane protein that is packaged into virions (65), but was not previously recognized to bind RNA. We noticed that ORF7a was significantly enriched in SHIFTR experiments targeting both the 5ʹ and 3ʹ terminal regions of the viral genome, but not when targeting the ORF1ab or sgmRNA regions (Figure 4A, Table S5). This binding pattern was unique among viral proteins and may hint at functions in genome cyclisation or viral RNA packaging that have been described for the terminal regions of SARS-CoV-2 (66, 67) and warrant further investigation. Together, these data demonstrate that SHIFTR is a powerful tool for dissecting functional interactions of distinct RNA regions and sequence elements within larger RNA molecules in intact cells. Using SHIFTR, we recover known and novel interactions of different sequence regions in the SARS-CoV-2 genome and reveal binding patterns of host and viral proteins that are consistent with the functions of their target RNA regions.

### Mapping of direct protein binding sites using eCLIP confirms binding preferences captured with SHIFTR

To further corroborate interactions identified with SHIFTR, we used eCLIP to map the binding pattern of several selected candidate proteins of host and virus with nucleotide resolution. As expected, eCLIP confirmed direct binding of all tested candidates (CNBP (19), LARP4, NSP9, ZAP) to SARS-CoV-2 RNA (Figure 5A). Moreover, the binding preferences observed by eCLIP are in line with the specific enrichment of candidate RBPs in SHIFTR experiments targeting defined sequence regions in the SARS-CoV-2 genome (Figure 5A). Using our previously published data (19), we confirmed substantial binding of CNBP to the region encoding all sgmRNAs in the SARS-CoV-2 genome, which matches the enrichment of CNBP in SHIFTR experiments (Table S5). For the host RNA-binding protein LARP4, we observed an enrichment of eCLIP binding sites above background mostly in regions encoding sgmRNAs, but also detected signal near the 5ʹ leader as well as the 3ʹ UTR (Figure 5A). As noted earlier, both sequences are present in all sgmRNAs. This binding profile is consistent with the strong enrichment of LARP4 in SHIFTR experiments targeting viral sgmRNAs and the 3ʹ UTR region (Table S5). Most notably, binding of the viral protein NSP9 to the 5ʹ leader sequence (Figure 4A) was fully corroborated by eCLIP, as indicated by a significantly enriched binding site at the 5ʹ end of the leader sequence (Figures 5A-B). Finally, ZAP, an antiviral protein induced by interferon that represses SARS-CoV-2 frameshifting (55), displayed widespread binding across ORF1ab and we identified a significantly enriched binding site overlapping the SARS-CoV-2 FSE (Figure 5C). However, we also noted that the ZAP binding profile extends to the region where sgmRNAs are produced (Figure 5A), which is again consistent with the enrichment of ZAP in SHIFTR experiments targeting this sequence region (Table S5). ZAP may execute its antiviral properties via multiple mechanisms and modes of binding that may involve destabilization of directly bound viral RNAs in addition to regulating frameshifting, as previously proposed (68, 69).

**Figure 5:**
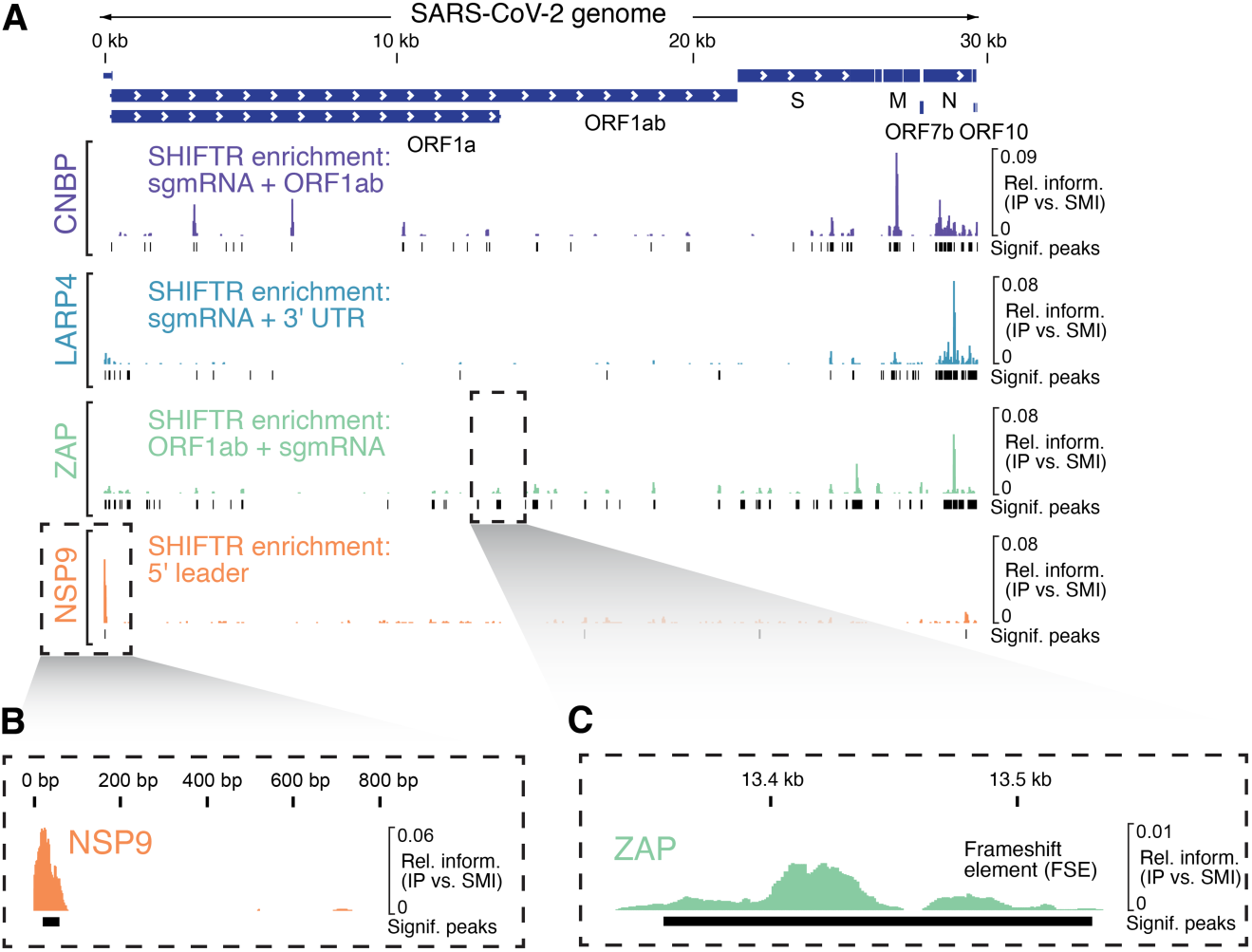
eCLIP validates binding preferences uncovered with SHIFTR. **(A)** Alignment of eCLIP data for CNBP (19) (human), LARP4 (human), NSP9 (SARS-CoV-2) and ZAP (human) to the SARS-CoV-2 RNA genome. Relative information in IP versus size-matched input (SMI) is calculated at each position and displayed along the viral genome. Protein binding sites significantly enriched relative to SMI are shown below coverage tracks. SHIFTR experiments displaying a significant enrichment of the target protein when interrogating different SARS-CoV-2 sequence regions are indicated in the order of the observed fold change next to coverage tracks. **(B)** As in **(A)**, but zoom-in to 5ʹ end as the dominant site of NSP9 binding is displayed. **(C)** As in **(A)**, but zoom-in region overlapping the frameshift element (FSE) is shown and ZAP binding profile is displayed.

Together, SHIFTR and eCLIP provide complementary evidence and help illuminate the distinct binding patterns and functions of host and viral protein on the SARS-CoV-2 RNA genome. We anticipate that SHIFTR will help prioritizing proteins for functional and mechanistic investigations based on their binding pattern. Moreover, the scalability and general applicability of the approach enables the systematic and region-resolved dissection of virtually all RNA regulatory elements in cellular transcriptomes in the future.

## DISCUSSION

Here, we present SHIFTR, a revolutionary tool enabling the region-resolved dissection of RNA-protein interactions for individual RNA elements in endogenous RNAs in an unbiased and highly scalable fashion. SHIFTR overcomes several roadblocks that previously limited the systematic characterization of RNA-protein interactions. First, SHIFTR reduces the amount of input material needed for the comprehensive characterization of RNA-protein interactomes by several orders of magnitude, which allows application of this technique to a large number of endogenously expressed RNAs in a systematic fashion and potentially using advanced model systems, such as primary cells or organoids. Second, the superior signal to noise ratio observed in SHIFTR experiments when compared to state-of-the-art RNA antisense purification-based technologies facilitates data interpretation and the selection of candidates for follow-up studies. Third, prior to the development of SHIFTR, it was not possible to identify interactomes for specific RNA regions within endogenous RNAs. Outside the context of an intact cell, *in vitro* approaches that combine synthetic RNA fragments with cell lysates in test tubes are commonly used (70). However, such approaches may not recapitulate interactions occurring in cells due to difficulties in controlling unspecific interactions and biases towards detecting highly abundant proteins (2). Hence, *in vitro* approaches are generally considered less well suited for interactome discovery. To investigate region-resolved RNA–protein interactions in the cellular environment, different technologies were developed to ectopically express RNA sequences of interest. In such an assay, the target RNA is frequently fused to an RNA aptamer sequence that can be harnessed to enrich the target RNA (71) or detect interactions with tagged RBPs using reporter-based overexpression systems (72). Alternative approaches, such as proximity-dependent protein labeling, rely on enzyme-catalyzed biotin labeling to identify proteins in proximity to target RNAs or RNA sequences (73). These approaches however require the establishment of genetically engineered cell systems and frequently do not differentiate direct from indirect interactors, which complicates the goal of identifying RNA region-specific interactions. SHIFTR overcomes these limitations and enables the region-resolved mapping of RNA-protein interactions for any endogenous RNA in any cell system without the need for genetic manipulation. Moreover, SHIFTR is a low-cost and easy to execute approach that requires no specialized equipment for the capture of direct RNA protein interactions. We provide streamlined workflows for targeting different RNA species that are heterogeneous in length and abundance, such as small non-coding RNAs, long non-coding RNAs, as well as mRNAs of virus and host. We show that SHIFTR is readily compatible with the sequential release of proteins bound to different RNAs of interest, thus enabling multiplexed RNA interactome capture in a single experiment. To facilitate the dissemination and application of SHIFTR, we provide Probe-SHIFTR, a computational tool for the design of optimized SHIFTR probe sets for any RNA type and sequence of interest, which includes species-specific filtering of probe sets in a fully customizable manner. Finally, we validate interactions and binding patterns observed with SHIFTR by eCLIP and observe large agreement in the recovered binding patterns, demonstrating the power of SHIFTR for revealing region-resolved RNA-protein interactions under endogenous conditions. In particular, we demonstrate that SHIFTR is ideally suited to investigate regulatory RNA elements in the genomes of RNA viruses such as SARS-CoV-2. Here, SHIFTR reveals distinct RNA binding preferences for known interactors in addition to uncovering previously unrecognized binding partners for specific sequence regions in the SARS-CoV-2 genome that are supported by functional data (58) and may point to novel host dependency mechanisms that could inspire the development of rationally designed therapeutic strategies. Among previously unrecognized SARS-CoV-2 RNA interactors discovered with SHIFTR is the regulatory subunit of the mRNA-capping methyltransferase RNMT:RAMAC complex. RAMAC is required for generating the 7-methylguanosine triphosphate (m^7^Gppp) moiety of the 5ʹ-cap of host mRNAs (74, 75). RAMAC has been proposed to be involved in cytoplasmic recapping of mRNAs that have lost their 5ʹ- cap and are translationally inactive and subject to decay (76, 77). Since the SARS-CoV-2 replication cycle crucially depends on the effective translation of both the RNA genome (ORF1ab) as well as the viral sgmRNAs, direct binding of RAMAC to both of these RNA species, as indicated by SHIFTR, may hint at the possibility of cytoplasmic recapping of viral mRNAs, which would maintain their translatability and impact viral gene expression. Beyond host proteins that bind SARS-CoV-2 RNAs, we also discovered viral proteins with intriguing binding patterns. First, SHIFTR accurately recovers binding of NSP9 to the 5ʹ-end of SARS-CoV-2 RNA, which is consistent with the emerging function of NSP9 in the biogenesis of SARS-CoV-2 RNAs at the priming or capping stage (60–62, 78). Second, ORF9b, a viral protein that is expressed from an alternative open reading frame (ORF) within the N gene, was significantly enriched in SHIFTR experiments targeting the sgmRNA and ORF1ab region. While ORF9b suppresses the host interferon response (79, 80), recent work suggests that ORF9b has a previously unrecognized RNA-binding activity (19, 20). Using SHIFTR, we uncover that the preferential binding of ORF9b to the ORF1ab and sgmRNA regions appears to mirror the binding pattern observed for N (Figure 4A), which may suggest that two RNA-binding proteins with a similar preference for binding viral RNA are expressed from the N gene.

In conclusion, with SHIFTR it is now possible to systematically and globally map interactions between the cellular proteome and any *cis*-regulatory RNA element from a diverse array of species. We envision that SHIFTR will revolutionize our understanding of site-specific RNA-protein interactions and regulatory events in complex transcriptomes, including pathogens and their hosts.

## CONTRIBUTIONS

M.M. conceived, designed, and supervised the study. J.A. developed SHIFTR and performed the majority of experiments. A.G. performed computational analyses and developed analytical concepts and ideas, including Probe-SHIFTR. S.Z. and N.S. helped with biochemistry and infection experiments, S.G. performed eCLIPs. S.A.C., M.S., and C.R.H. performed mass spectrometry work for RAP-MS. M.M. and J.A. wrote the manuscript with input from all authors.

## ACKNOWLEDGMENTS

We thank J. Vogel for continuous support. We are grateful to the EMBL Proteomic Core Facility, in particular J. Schwarz and F. Stein for conducting mass spectrometry experiments and supporting data analysis. We thank C. Krempl for overseeing infection experiments conducted by S.Z. and N.S. in BSL3 facility. We thank S. Pöhlmann for sharing Vero-E6-TMPRSS2 cells and A. Pichlmair for sharing A549^ACE2^ cells; the Core Unit Systems Medicine in Würzburg for sequencing; H. Keshishian and K. Clauser for help with MS database searches; A. Sparmann and the Munschauer group for helpful discussions and critical comments on the manuscript.

## MATERIALS AND METHODS

### Cell lines

Huh-7 cells (a generous gift from the Virology Diagnostics Unit at the Institute of Virology and Immunobiology, University of Würzburg), TMPRSS2-Vero-E6 cells (a generous gift from S. Pöhlmann), A549^ACE2^ cells (a generous gift from A. Pichlmair) and A549 cells were cultured in DMEM medium (31966047, Thermo Fisher Scientific) supplemented with 10 % (v/v) heat-inactivated FCS (10500064, Gibco), 100 U/ml streptomycin and 100 µg/ml penicillin (15140122, Gibco). Cells were maintained at 37 °C and 5 % CO_2_.

### Virus production and infection

Virus production and infection experiments were carried out as described in Schmidt *et al.*, (2020) (19). For SHIFTR and eCLIP experiments, we infected A549^ACE2^ cells with SARS-CoV-2 at MOI 5 PFU/cell. At 24 hpi, cells were UV crosslinked and harvested as described below.

### Design of SHIFTR DNA probes

DNA oligonucleotides were designed using the Probe-SHIFTR tool and synthesized as unmodified DNA oligonucleotides in plates or as oligo pools. For SARS-CoV-2 SHIFTR experiments we designed 50-mer oligonucleotides and synthesized probe sets targeting the ORF1ab and sgmRNA regions as oligo pools. For all other RNA targets, we designed 25-mer oligonucleotides. All probe sequences are provided in Supplementary Table S6. Probe-SHIFTR is a java workflow which creates all possible k-mers for a given RNA target sequence. Each k-mer is evaluated with respect to sequence features such as containing poly nucleotide chains, sequence complexity, repetitive sequences, and containing masked repeat regions. After filtering each k-mer based on these features, a similarity search is performed by BLAT to remove k-mers showing high similarity to genomic or transcriptomic regions of a provided reference specie. After performing these filtering steps each k-mer should represent a unique sequence of the target RNA region. Using a greedy algorithm, the resulting k-mers are arranged in sets of non-overlapping sequences providing maximal coverage over the target region while maintaining the unique sequence feature of this region. Probe-SHIFTR provides several sets of non-overlapping nucleotide sequences for different parameter settings as well as analysis plots helping to decide which set of oligonucleotides should be used. The software is publicly available at https://github.com/AlexGa/ProbeSHIFTR.

### Immunoblotting

Immunoblots were performed as described previously (19) using the following antibodies: anti-SNRNP70 (ab83306, abcam) and anti-HEXIM1 (15676-1-AP, Proteintech).

### Silver staining

Silver staining was performed using Pierce’s Silver Stain Kit according to the manufacturer’s instructions (24612, Thermo Fisher Scientific).

### *In vitro* transcription of U1 RNA

The DNA template was generated by cloning the U1-1 sequence (NR_004430.2) into the pCR4-TOPO backbone (Thermo Fisher Scientific). Plasmid DNA was linearized with NotI-HF (R3189, NEB) and subjected to IVT reactions using 50 ng/μl T7 RNA polymerase HC (EP0113, Jena Bioscience), with equimolar ratios of all natural ribonucleotides. The complete IVT mixture was incubated for 2 h at 37 °C. To remove any residual DNA template, 50 U/ml DNase I (M0303, NEB) were added and samples were incubated for another 30 min at 37 °C. RNA was purified using the RNA Clean and Concentrator-5 kit (R1013, Zymo Research).

### SHIFTR

#### Crosslinking and pre-clearing

For each target RNA 1×10^7^ Huh-7 or A549 cells were used per replicate. Cells were washed twice with cold PBS and crosslinked in a GS Gene Linker (Bio-Rad Laboratories) using 0.8 J/cm^2^ of 254 nm UV light. The crosslinked cells were scraped off in cold PBS and lysed in 281 μl cold lysis buffer (50 mM Tris-HCl pH 7.5, 150 mM NaCl, 1 % IGEPAL (NP-40), 0.5 % sodium deoxycholate, 0.5 % n-Dodecyl ß-maltoside (DDM), 5 mM DTT, 1x complete protease inhibitor cocktail (A32955, Thermo Fisher Scientific) and 100 U/ml murine RNase inhibitor (M0314, NEB)) for 20 minutes on ice, resuspending every 5 minutes. Next, 9 μl of a 100 mM MgCl_2_ stock and 10 μl of 2 U/μl TURBO DNase (AM2238, Thermo Fisher Scientific) were added. Lysates were incubated for 30 min at 37 °C with occasional mixing by inversion. After centrifugation for 10 min at 14, 000 g and 4 °C, 300 μl of the supernatant were transferred to a new tube and mixed with 3 volumes (900 μl) of Trizol LS (T3934, Sigma-Aldrich) by vortexing. After incubation for 5 min at room temperature, lysates were frozen at -80 °C for up to 4 weeks or used immediately. For the experiment omitting the pre-clearing step, cells were directly lysed in 1 ml of Trizol (R2050-1, Zymo Research) and subjected to interface cleanup as described below.

#### Multiplexed poly(A)-SHIFTR

For the multiplexed identification of RNA interactomes for moderately abundant cellular mRNAs and lncRNAs, we subjected 1.5×10^8^ cells to UV crosslinking as described above. To enrich polyadenylated RNA, oligo(dT) beads (NEB, S1419S) were used. Cell pellets were lysed in 10:1 (w/w) of lysis buffer (100 mM Tris-HCl pH 7.5, 500 mM LiCl, 0.5 % LDS, 1 mM EDTA, 5 mM DTT, 1x complete protease inhibitor cocktail, 100 U/ml murine RNase inhibitor), passed 5x through a 0.8 mm gauge needle and once through a 0.4 mm gauge needle. Lysates were centrifuged for 15 min at 5, 000 g and 4 °C and supernatants were transferred to new 50 ml tubes. For each sample 3 ml of oligo(dT) beads were pre-conditioned with 3 ml of lysis buffer by mixing and rotating for 5 min, before removing the supernatant on a magnet. The beads were added to the lysates and samples were incubated 1 h at room temperature with gentle rotation. After separation on the magnet, supernatants were transferred to new tubes and saved for another round of poly(A) capture. Beads were washed twice using Wash Buffer 1 (20 mM Tris-HCL pH 7.5, 500 mM LiCl, 0.1 % LDS, 1 mM EDTA, 5 mM DTT, 1x protease inhibitor), twice using Wash Buffer 2 (20 mM Tris-HCl pH 7.5, 500 mM LiCl, 0.02 % NP-40, 1 mM EDTA, 5 mM DTT, 1x protease inhibitor) and once using Wash Buffer 3 (20 mM Tris-HCl pH 7.5, 200 mM LiCl, 0.02 % NP-40, 1 mM EDTA, 5 mM DTT, 1x protease inhibitor). To elute the bound RNA-protein complexes, beads were resuspended in 180 μl Elution Buffer (20 mM Tris-HCl pH 7.5, 1 mM EDTA) and transferred into 5 ml tubes. For heat elution, samples were incubated 5 min at 80 °C, before placing them on a magnet and quickly transferring supernatants into 1.5 ml tubes. The poly(A) capture was repeated twice, using the saved lysates. The pooled eluate with a volume of ∼600 μl was split into two 1.5 ml tubes and 900 μl Trizol LS were added to each tube. After the first phase separation and removal of aqueous and organic phases (see “Interface cleanup”), the two interfaces were resuspended in a total of 1 ml Trizol and combined into a single 1.5 ml tube. Interface cleanup and subsequent RNase H digest were performed as described below.

#### Interface cleanup

To each sample 220 μl chloroform were added and samples were mixed by vortexing. After phase separation by centrifugation for 15 min at 12, 000 g and 4 °C, most of the upper aqueous and lower organic phase were removed (< 100 µl remaining). If necessary, centrifugation was repeated to remove additional volume without losing interface material. 1 ml Trizol (R2050-1, Zymo Research) as well as 200 µl chloroform were added to the interface and phase separation and removal were repeated. After a total of 4 rounds of phase separation, 9 volumes of methanol were added to the final interface for precipitation as previously described (10). After vortexing and centrifugation for 10 min at 14, 000 g and 4 °C, all supernatant was removed and the precipitate was washed once with 1 ml methanol. Without drying the pellet, 126 μl H_2_O (111 μl for multiplexed poly(A)-SHIFTR) were added and samples were incubated for 1 h on ice to reconstitute interfaces.

#### RNase H digest

After thoroughly resuspending the interface precipitate, 150 μl 2x RNase H digestion buffer (100 mM Tris-HCl pH 8.3, 150 mM KCl, 6 mM MgCl_2_, 20 mM DTT, 1 % (v/v) NP-40, 1 % (v/v) Triton X-100, 1 % (v/v) DDM) were added. Next, we added 15 μl (30 μl for multiplexed poly(A)-SHIFTR) of each 100 μM SHIFTR probe set. At this point, after a short spin 10 μl of the reconstituted interface were separated and set aside for RNA analysis. Next, 9 μl of 5 U/μl of thermostable RNase H (M0523, NEB) were added and samples were incubated 1 h at 50 °C. After target RNA digest and another short spin, we separated 10 μl of the reconstituted interface for RNA analysis. Next, 3 volumes (840 μl) of Trizol LS were added and samples mixed by vortexing.

For the experiment involving Xrn1 exonuclease digestion after RNase H treatment, 2 μl 1M MgCl_2_ and 2 μl of 1 U/μl Xrn1 (M0338, NEB) were added after RNase H digest. Samples were incubated for 1 h at 37 °C, before Trizol LS was added as described above.

#### Protein extraction

To each sample 200 μl chloroform were added and samples were mixed by vortexing. After centrifugation for 15 min at 12, 000 g, and 4 °C, the aqueous phase was removed while the interphase was transferred to a new tube and set aside*. About 75 % of the remaining organic phase were cleanly transferred to a new tube for protein precipitation. Next, 130 µl of each organic phase were transferred to new tubes and mixed with 9 volumes (1170 µl) of methanol by vortexing. After centrifugation for 10 min at maximal speed and 4 °C, supernatants were discarded and another 130 µl of organic phases were added and precipitated following the described procedure. This was repeated until most of the organic phases had been collected, using equal volumes for all samples. Pellets were finally washed once with 1 ml methanol and briefly air-dried.

*For multiplexed poly(A) SHIFTR, which requires multiple sequential RNase H digestion steps, interphases set aside during protein extraction were re-dissolved in 1 ml Trizol and stored at - 80 °C or immediately mixed with 200 μl chloroform for another round of phase separation. After centrifugation for 15 min at 12, 000 g and 4°C, the aqeuous phases and organic phases were removed. Interfaces were precipitated using methanol as described above and reconstituted for another round of RNase H digest.

### RNA antisense capture from interfaces

For RNA antisense capture, precipitated interfaces were taken up in 300 μl H_2_O and incubated 1 h on ice. After thoroughly resuspending the interface, 600 μl of 1.5x hybridization buffer (15 mM Tris pH 7.4, 7.5 mM EDTA, 750 mM LiCl, 0.75 % DDM, 0.3 % SDS, 0.15 % NaDC, 6 M urea, 7.5 mM DTT) were added and 10 μl were removed for RT-qPCR analysis. For each sample 1.5 μg of biotinylated DNA antisense probes (1 μg/μl) were denatured for 3 min at 85 °C, placed on ice and subsequently added to the samples. GFP targeting probes were used for control samples. Hybridization was performed for 2 h at 50 °C with 900 rpm shaking (15 s on/15 s off). For each sample 150 μl Dynabeads MyOne Streptavidin C1 (65001, Thermo Fisher Scientific) were washed four times using 150 μl 10 mM Tris pH 7.5 and twice using 1x hybridization buffer (10 mM Tris pH 7.4, 5 mM EDTA, 500 mM LiCl, 0.5 % DDM, 0.2 % SDS, 0.1 % NaDC, 4 M urea, 5 mM DTT). Using 100 μl of each sample, beads were resuspended and transferred to the sample. For bead capture, samples were incubated 45 min at 50 °C with 900 rpm shaking (15 s on/15 s off). After bead separation on a magnet, supernatants were transferred to new tubes and stored at -80 °C. Beads were washed three times using each 1 ml pre-heated 1x hybridization buffer, each time removing supernatants on magnet, resuspending beads in fresh buffer, transferring the suspensions to new 2 ml tubes and incubating 5 min at 50 °C with 800 rpm shaking (15 s on/15 s off). Before elution, beads were rinsed on-magnet with 500 μl RNase H digest buffer and subsequently resuspended in 301 μl of RNase H digest buffer. 10 μl of the bead suspension were removed for RT-qPCR analysis, before 9 μl of 5 U/μl thermostable RNase H (M0523, NEB) were added and samples were incubated 1 h at 50 °C with 800 rpm shaking (15 s on/15 s off). After separation on magnet, supernatants were transferred to 1.5 ml tubes. Another 100 μl of buffer were added to beads, mixed and supernatants were combined after separation on magnet. The combined supernatant (∼400 μl) was transferred to new tubes two additional times after incubation on magnet to remove any residual beads. For protein precipitation, 100 μl of 100 % TCA were added to a final concentration of 20 % TCA. After incubation overnight covered by ice at -20 °C, samples were centrifuged 1 h at 15, 000 g and 4 °C. Supernatants were transferred to new tubes and pellets were washed with 100 μl ice-cold acetone, before centrifugation for 30 min at 15, 000 g and 4 °C. Supernatants were removed and samples were briefly air-dried.

### RNA extraction

10 μl of the reconstituted interface (see SHIFTR section) or antisense capture samples were briefly denatured at 95 °C and treated with 28 μl Proteinase K Buffer (20 mM Tris pH 8.5, 10 mM EDTA, 2 % NLS, 2.5 mM TCEP) as well as 2.5 μl of 0.8 U/µl proteinase K (NEB). After protein digestion for 90 minutes at 55 °C and 800 rpm (15s on/15 s off), 30 μl of the samples were cleaned using MyOne Silane Dynabeads (Thermo Fisher Scientific). For each sample, 15 μl beads were washed twice in RLT Buffer and applied to sample in 3 sample volumes (90 μl) of RLT Buffer. Upon addition of 4.5 volumes (135 μl) of isopropanol, samples were incubated for 10 min at room temperature and washed twice with 70 % ethanol. Beads were air-dried and resuspended in 30 μl DNase digestion mix (1x TURBO DNase buffer, 1 U/μl murine RNase inhibitor and 0.2 U/μl TURBO DNase). After incubation for 30 min at 37 °C, another Silane bead cleanup was performed and RNA was eluted in 18 μl H_2_O. 8 μl of the RNA were used for either RT-qPCR or RNA-seq analysis.

### RT-qPCR

For reverse transcription, 1 μl random primer mix (9-mer, 100 μM) was added to 8 μl of sample and denatured for 3 min at 70 °C. Primers were annealed for 10 min at RT. 11 μl of RT master mix (2 μl 10x Affinity Script buffer, 2 μl 100 mM DTT, 1 μl Affinity Script Enzyme, 0.8 μl 100 mM dNTPs) were added and samples were incubated 1 h at 55 °C, followed by inactivation for 15 min at 70 °C. For RT-qPCR, samples were diluted 1:5 or 1:100 (for 18 S rRNA measurement). 12.75 μl of the diluted cDNA were mixed with 14.25 μl qPCR master mix (13.5 μl Luna Universal qPCR Master Mix (NEB) and 0.75 μl of 25 μM primer mix) and measured in technical quadruplicates using the Quant Studio 5 system. RT-qPCR primer sequences are listed in Supplementary Table S6.

### Mass spectrometry

#### Sample Preparation

Reduction of disulfide bonds on cysteine was performed with dithiothreitol (56°C, 30 min, 10 mM in 50 mM HEPES, pH 8.5) followed by alkylation with 2-chloroacetamide (room temperature, in the dark, 30 min, 20 mM in 50 mM HEPES, pH 8.5). The SP3 protocol (26, 27) used for sample clean-up and trypsin (sequencing grade, Promega) was added in an enzyme to protein ratio 1:50 for overnight digestion at 37°C (in 50 mM HEPES).

#### TMT Labeling

Peptides were labelled either with TMT6plex (28) (experiments P2490 (poly(A)-SHIFTR ACTIN (elution 1)) and P2604 (poly(A)-SHIFTR MALAT1 (elution 2) & GAPDH)), TMT10plex (29) (experiments P1883 (SHIFTR AGPC only), P1901 (SHIFTR Pre-clear + AGPC), P2349 (U1 SHIFTR A549 deep MS) and P2664 (SARS-CoV-2 SHIFTR)) or TMT16plex (30) (experiments P2107 (SHIFTR RNase H comparison) and P2508 (SHIFTR RAP comparison) Isobaric Label Reagent (Thermo Fisher) according the manufacturer’s instructions. In brief, of 0.8 mg reagent dissolved in 42 µl acetonitrile (100 %) 8 μl was added and incubated for 1 h at room temperature. The reaction was stopped with 8 μl 5 % hydroxylamine and incubated for 15 min at room temperature. Samples of a set were combined and desalted on an OASIS HLB µElution Plate (Waters). Offline high pH reverse phase fractionation (experiments P2107 (SHIFTR RNase H comparison), P2349 (U1 SHIFTR A549 deep MS), P2490 (poly(A)-SHIFTR ACTIN (elution 1), P2508 (SHIFTR RAP comparison), P2604 (poly(A)-SHIFTR MALAT1 (elution 2) & GAPDH), P2664 (SARS-CoV-2 SHIFTR)) was carried out on an Agilent 1200 Infinity high-performance liquid chromatography system, equipped with a Gemini C18 column (3 μm, 110 Å, 100 x 1.0 mm, Phenomenex) installed with a Gemini C18, 4 x 2.0 mm SecurityGuard (Phenomenex) cartridge as a guard column. The binary solvent system consisted of 20 mM ammonium formate (pH 10.0) (A) and 100 % acetonitrile as mobile phase (B). The flow rate was set to 0.1 mL/min. Peptides were separated using a gradient of 100 % A for 2 min, to 35 % B in 59 min, to 85 % B in another 1 min and kept at 85 % B for an additional 15 min, before returning to 100 % A and re-equilibration for 13 min. 48 fractions were collected which were pooled into 6 fractions. Pooled fractions were dried under vacuum centrifugation, reconstituted in 10 μl 1 % formic acid, 4 % acetonitrile and then stored at -80 °C until LC-MS analysis.

#### Data Acquisition

An UltiMate 3000 RSLC nano LC system (Dionex) fitted with a trapping cartridge (µ-Precolumn C18 PepMap 100, 5 µm, 300 µm i.d. x 5 mm, 100 Å) and an analytical column (nanoEase™ M/Z HSS T3 column 75 µm x 250 mm C18, 1.8 µm, 100 Å, Waters) was coupled to an Orbitrap Fusion Lumos Tribrid Mass Spectrometer (Thermo) using the Nanospray Flex ion source in positive ion mode. The samples were applied onto the trapping column with a constant flow of 30 µL/min 0.05 % trifluoroacetic acid in water for 6 minutes. After switching in line with the analytical column peptides were eluted at a constant flow of 0.3 µL/min using the method described in the following. The binary solvent system consisted of 0.1 % formic acid in water with 3 % DMSO (solvent A) and 0.1 % formic acid in acetonitrile with 3 % DMSO (solvent B). For unfractionated samples a 120 min method was used (experiments P1883 (SHIFTR AGPC only), P1901 (SHIFTR Pre-clear + AGPC), P2107 (SHIFTR RNase H comparison)): The percentage of solvent B was increased from 2 % to 8 % in 4 min, from 8 % to 28 % in 104 min, from 28 % to 40 % in another 4 min and finally from 40 % to 80 % in 4 min, followed by re-equilibration back to 2 % B in 4 min.

For fractionated samples either a 60 min method in which solvent B was increased from 2 % to 8 % in 6 min, then from 8 % to 28 % in 42 min (for TMT16 to 30 % in 36 min), from 28 % to 40 % in 4 min (for TMT16 from 30 % to 40 % in 10 min) from 40 to 80 % in 4 min (for TMT16 in 3 min) and a re-equilibration to 2 % B for 4 min (for TMT16 in 5 min) (experiments P2490 (poly(A)-SHIFTR ACTIN (elution 1), P2508 (SHIFTR RAP comparison), P2604 (poly(A)-SHIFTR MALAT1 (elution 2) & GAPDH)) or a 90 min method was used in which solvent B was raised from 2 % to 8 % in 6 min, from 8 % to 28 % in 72 min (for TMT16 to 30 % in 66 min), from 28 % to 38 % in 4 min (for TMT16 to 40 % in 10 min), from 38 % to 80 % in 4 min (for TMT16 to 80 % in 3 min) and re-equilibration back to 2 % B for 4 min (for TMT16 5 min) (experiments P2107 (SHIFTR RNase H comparison), P2349 (U1 SHIFTR A549 deep MS), P2664 (SARS-CoV-2 SHIFTR)).

The peptides were introduced into the MS instrument via a Pico-Tip Emitter 360 µm OD x 20 µm ID; 10 µm tip (New Objective) and an applied spray voltage of 2.4 kV. The capillary temperature was set at 275°C. Full mass scan was acquired with mass range 375-1500 m/z in profile mode in the orbitrap with resolution of 60000 (TMT6) or 120000 (TMT10 and TMT16). The filling time was set at maximum of 50 ms with a limitation of 4x10^5^ ions. Data dependent acquisition (DDA) was performed with the resolution of the Orbitrap set to 15000 (TMT6) or 30000 (TMT10 and TMT16), with a fill time of 54 (TMT6) or 94 ms (TMT10 and TMT16) and a limitation of 1x10^5^ ions. A normalized collision energy of 34 (TMT16) or 36 (TMT6 and10) was applied. MS^2^ data was acquired in profile mode. Define first mass was set to 110 m/z.

#### Quantification and identification of peptides and proteins

IsobarQuant (31) with Mascot (v2.2.07) (P1883 (SHIFTR AGPC only), P1901 (SHIFTR Pre-clear + AGPC), P2107 (SHIFTR RNase H comparison), P2349 (U1 SHIFTR A549 deep MS)) or MS Fragger v3.7 (32) (experiments P2490 (poly(A)-SHIFTR ACTIN (elution 1), P2508 (SHIFTR RAP comparison), P2604 (poly(A)-SHIFTR MALAT1 (elution 2) & GAPDH) and P2664 (SARS-CoV-2 SHIFTR)) was used to process the acquired data, which was searched against a Homo sapiens proteome database (IsobarQuant: UP000005640, May 2016, 92507 entries; MSFragger: UP000005640, October 2022, 20594 entries) plus common contaminants and reversed sequences. For experiment P2664 (SARS-CoV-2 SHIFTR) also the SARS-CoV-2 database was used (UP000464024, February 2023, 31 entries). The following modifications were included into the search parameters: Carbamidomethyl on Cysteine and TMT6/10/16 on lysine as fixed modifications, protein N-term acetylation, oxidation on methionine and TMT6/10/16 on N-termini as variable modifications. For precursor ions a mass error tolerance of 10 ppm (IsobarQuant) or 20 ppm (MSFragger) was used and for fragment ions 0.02 Da (IsobarQuant) or 20 ppm (MSFragger) was set. Trypsin was set as protease with a maximum of two missed cleavages. The minimum peptide length was set to seven amino acids. At least two unique peptides were required for a protein identification. The false discovery rate on peptide and protein level was set to 0.01.

#### Statistical data analysis

The raw output files of IsobarQuant (protein.txt – files) were processed using the R programming language. Only proteins that were quantified with at least two unique peptides were considered for the analysis. Raw TMT reporter ion signals (signal_sum columns) were first cleaned for batch effects using limma (33) and further normalized using vsn (variance stabilization normalization (34). Proteins were tested for differential expression using the empirical Bayes statistics (eBayes) from the limma package while removing batch effects by incorporating replicate information. For correcting against multiple testing, the local false discovery rate (FDR) was calculated by fdrtool (35). Comparisons showing a local FDR value below 0.05, were considered as statistically significant. As described in Strimmer et al. (35), the local FDR determines an upper bound for tail-based FDR values provided by fdrtool. To facilitate the identification of additional candidates in experiments for MALAT1, GAPDH, and ACTB, the tail-based FDR value (qvalue) was considered for differentiating between statistically significant and non-significant comparisons.

Comparing the performance of SHIFTR and RAP-MS for U1 and 7SK, both approaches were regarded as binary classifiers distinguishing target proteins (positives) by having low FDR values and background noise (negatives) having high FDR values. Based on the detailed knowledge about the composition of both snRNPs, the number of true positives (TP), false positives (FP), true negatives (TN), and false negatives (FN) could be calculated (see table below). Based on this definition, we calculated precision as TP/(TP + FP), recall as TP/(TP + FN), and the F1-score as the harmonic mean of both measures as 2 × (precision × recall)/(precision + recall). Additionally, using the negative logarithm of FDR values as classification scores, the calculations for precision-recall curves could be performed by using the R package PRROC (36). These curves serve as an indication how precisely each method is able to distinguish target proteins from background noise.

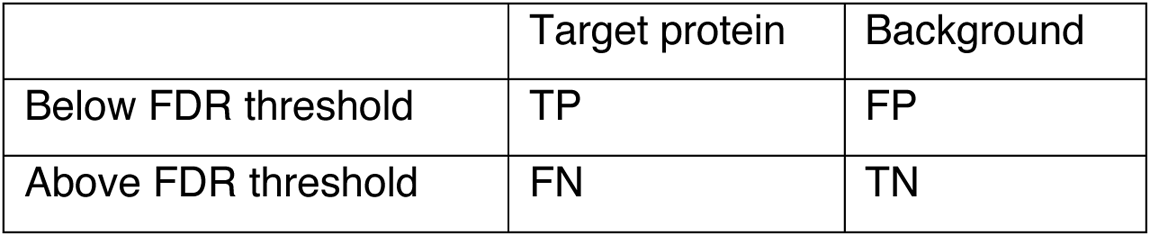

|  | Target protein | Background |
| --- | --- | --- |
| Below FDR threshold | TP | FP |
| Above FDR threshold | FN | TN |

### RNA antisense purification and mass spectrometry (RAP-MS)

RAP-MS, including mass spectrometry sample preparation was performed as described in Munschauer *et al.,* (2018) (16).

### Enhanced crosslinking and immunoprecipitation (eCLIP)

We performed eCLIP for selected RNA-binding proteins (LARP4, ZAP, NSP9) in SARS-CoV-2 infected A549^ACE2^ cells as described previously in Schmidt et al., (2020) (19). The following antibodies were used: LARP4: Proteintech, 16529-1-AP; ZAP: Proteintech, 16820-1-AP, NSP9: Genetex, GTX135732. Resulting libraries were sequenced paired-end with a read length of 2 x 40 nucleotides. Paired-end sequencing reads were adapter- and quality trimmed using cutadapt (v1.18). Reads with a total length less than 18 nt were discarded. A custom java program was applied that simultaneously identified and clipped the remaining unique molecular identifier (UMI) associated with each read. These trimmed reads were then aligned to the human (hg38, Ensembl release 106) and the SARS-CoV-2 reference genome (NC_045512.2, GenBank: MN908947.3) using STAR (v2.7.10a) (37) with the parameters – outFilterScoreMinOverLread 0 –outFilterMatchNminOverLread 0 –outFilterMatchNmin 0 – outFilterType Normal –alignSoftClipAtReferenceEnds No --alignSJoverhangMin 8 – alignSJDBoverhangMin 1 –outFilterMismatchNoverLmax 0.04 --scoreDelOpen -1 – alignIntronMin 20 --alignIntronMax 3000 –alignMatesGapMax 3000 –alignEndsType EndToEnd. Next, we removed PCR duplicates using the UMI-aware deduplication functionality in Picard’s MarkDuplicates. Regions with enriched protein binding were identified with MACS2 (38) by modeling the fold change in IP samples over a paired SMI control using the parameters -g 29903 -s 31 --keep-dup all --nomodel --d-min 25 --call-summits --scale-to small --shift 25 --nolambda --extsize 5 --max-gap 20 --min-length 5. The identified MACS2 peaks were additionally filtered by calculating the enrichment of reads within each peak over all remaining mapped reads between IP and size matched input. A statistically significant enrichment relative to SMI control was calculated by a one-sided Fisher’s exact test. The resulting *P* values were corrected with the Benjamini-Yekutieli (39) procedure and only peaks with an adjusted *P* value < 0.05 were considered for further analysis.

The overall eCLIP signal was visualized by calculating the relative information content of IP over SMI (40–42). The relative information content was defined as *p_i_* x *log_2_(p_i_/q_i_)*, where *i* denotes a certain genomic position, *p_i_* represents the fraction of total aligned reads in IP and *q_i_* represents the fraction of total aligned reads in SMI. Visualizations of the region were rendered from the PCR-deduplicated .bam files using the Integrative Genome Visualization (IGV) Browser.

### RNA sequencing

For removal of ribosomal RNA, the NEBnext rRNA depletion kit v2 (E7400) was used according to the manufacturer’s instructions and samples were eluted in 9 μl H_2_O. Next, 1 μl 10x FastAP buffer (Thermo Fisher) was added and RNA was fragmented for 3 min at 90 °C, before placing samples on ice. After addition of 2.5 μl FastAP master mix (1x FastAP buffer, 4 U/μl murine RNase inhibitor, 0.7 U/μl of FastAP enzyme (EF0651, Thermo Fisher Scientific), dephosphorylation was performed 10 min at 37 °C. Next, 17.5 μl of T4 PNK master mix (1.7x T4 PNK Buffer (NEB), 570 U/μl murine RNase inhibitor, 28 mU/μl TURBO DNase, 0.8 U/μl T4 PNK (NEB)) were added and samples incubated another 20 min at 37 °C. Next, Silane bead cleanup was performed as described previously (see section “RNA extraction”) and RNA was eluted in 7 μl H_2_O. For poly-A-tailing, 3 μl of cold Poly-A-tailing master mix (3.3x Smartscribe first strand buffer (Takara), 0.16 U/μl Poly-A-polymerase (2180A, Takara), 3.3 U/μl SuperaseIn RNase inhibitor (AM 2694, Thermo Fisher Scientific), 0.33 mM ATP (N0437A, NEB)) were prepared and added on ice and samples were incubated 10 min at 37 °C, before inactivating 20 min at 65 °C. For reverse transcription, 1 μl of oligo dT primer was added and annealed by incubating 3 min at 72 °C before gradually cooling to 4 °C over a span of 2 minutes. 8 μl of Smart RT master mix (1.25x Smartscribe first strand buffer, 0.625 U/μl SuperaseIn RNase inhibitor, 12.5 U/μl Smartscribe reverse transcriptase (639538, Takara), 2.5 mM dNTPs, 1.25 mM DTT) were added and reverse transcription was performed 15 min at 42 °C. Template switching oligo (1 μl of a 10 μM stock) was added to a final concentration of 0.5 μM and RT reaction performed for another 90 min at 42 °C. After inactivation for 10 min at 70 °C, cDNA was diluted with 55 µl H_2_O and 21 µl of the diluted cDNA were used for PCR amplification. Next, 29 μl of PCR master mix (2x Q5 high-fidelity master mix (NEB), 1 μM 2P universal forward primer, 1 μM barcoded reverse primer) were added and libraries were amplified for 12-14 cycles. 2 μl of the PCR reactions were separated on a 2 % agarose gel to assess amplification. For library purification, 1 volume of AMPure XP beads (A63881, Beckmann Coulter) was added and samples incubated for 10 min at room temperature. Beads were washed twice using 80 % ethanol, air-dried and libraries eluted in 20 μl H_2_O. After separation on a 2 % agarose gel, gel fragments corresponding to 190 – 350 nts in DNA size were excised and libraries extracted using the Zymoclean Gel DNA Recovery Kit (D4007, Zymo Research). DNA concentrations were measured using the Qubit dsDNA HS Assay Kit (Q32854, Life Technologies) and cDNA libraries were sequenced using the Illumina NextSeq platform. Sequences of oligonucleotides used for RNA sequencing library preparation are listed in Supplementary Table S6.

### Gene Ontology (GO) enrichment analysis

We performed Gene Ontology (GO) enrichment analyses for the various SARS-CoV-2 RNA interactomes and protein subgroups specified in the main text using the Database for Annotation, Visualization and Integrated Discovery (DAVID) (43) tool (https://david.ncifcrf.gov/tools.jsp) and applying default settings.

## SUPPLEMENTARY FIGURES

**Figure S1:**
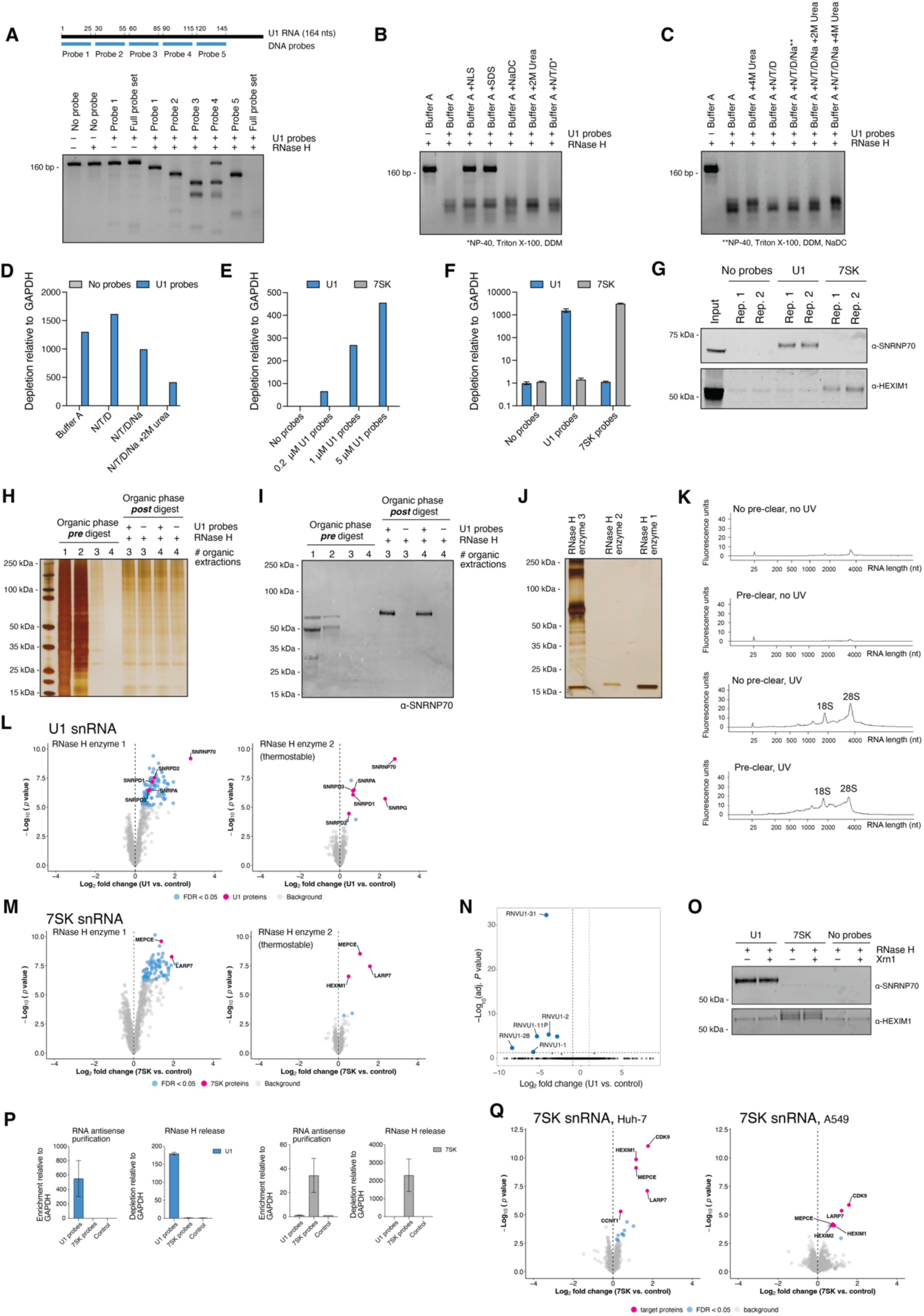
Optimization of experimental parameters for the effective digestion of target RNAs and the release of bound proteins from crosslinked interfaces. **(A)** Gel electrophoresis analysis of *in vitro* transcribed U1 RNA and the cleavage patterns observed with DNA probes targeting different RNA regions as illustrated in schematic shown on top. **(B)** Gel electrophoresis analysis of *in vitro* transcribed U1 RNA and its digestion by RNase H using sequence-specific DNA probes in buffers supplemented with different detergents and chaotropic agents as indicated. Buffer A: 50 mM Tris-HCl, 75 mM KCl, 3 mM MgCl_2_, 10 mM DTT. **(C)** As in **(B)**, but supplementing digestion buffer with additional detergents and/or chaotropic agents. **(D)** RT-qPCR analysis of RNase H-mediated depletion of endogenous U1 RNA in digestion buffers supplemented with different detergents and chaotropic agents analyzed in (A) and (B). Interfaces from UV-crosslinked Huh-7 cells were used. Depletion is normalized to GAPDH and compared to samples treated with RNase H in the absence of DNA probes (No probes). **(E)** RT-qPCR analysis of RNase H-mediated depletion of endogenous U1 RNA using different concentrations of DNA probes. Interfaces from UV-crosslinked Huh-7 cells were used. Depletion is normalized relative to GAPDH. Unspecific cleavage is estimated by measuring changes to 7SK RNA levels. **(F)** RT-qPCR analysis of RNase H-mediated depletion of the endogenous U1 or 7SK snRNA in interfaces isolated from UV-crosslinked cells. Depletion is normalized relative to GAPDH. Values are mean ± standard error of the mean. **(G)** Western blot analysis of proteins released from UV-crosslinked interfaces after the targeted digestion of U1 or 7SK snRNAs using sequence-specific DNA probes together with RNase H. SNRNP70 and HEXIM1 serve as positive controls for the U1 and 7SK snRNP complexes, respectively. Two independent replicates were analyzed. **(H)** Silver staining of SDS polyacrylamide gel analyzing the protein content of organic phases after several consecutive phase extraction steps before and after RNase H treatment of interfaces with and without U1-specific DNA probes. **(I)** Western blot analysis of protein content of organic phases after several consecutive phase extraction steps before and after RNase H treatment of interfaces with and without U1-specific DNA probes. SNRNP70 serves as positive controls for the release of U1 snRNP complex components. **(J)** Silver staining of SDS polyacrylamide gel analyzing the protein content of several different commercially available RNase H enzyme preparations. **(K)** Electropherogram of capillary electrophoresis analysis of RNA isolated from UV-crosslinked or uncrosslinked interfaces with and without pre-clearing after protein removal by Proteinase K treatment. **(L)** Quantitative comparison of proteins released from UV-crosslinked interfaces when using different commercially available RNase H enzymes for SHIFTR experiments targeting the U1 snRNA. SHIFTR was performed with pre-clearing. Volcano plots display log_2_ fold changes comparing U1-depleted to untreated samples. Known components of the U1 snRNP complex are highlighted in pink. Mass spectrometry experiments were performed without offline fractionation (Methods). **(M)** As in **(L)**, but for the 7SK snRNA. Known components of the 7SK snRNP complex are highlighted in pink. **(N)** Differential gene expression analysis of UV-crosslinked interfaces after U1 degradation with thermostable RNase H and sequence-specific DNA probes compared to interfaces subjected to treatment without the addition of probes and enzyme (control). Differentially expressed genes are highlighted in blue. **(O)** Western blot analysis of proteins released from UV-crosslinked interfaces after the targeted digestion of U1 or 7SK snRNAs using sequence-specific DNA probes together with RNase H. RNase H treatment was followed with a 1 h Xrn1 exoribonuclease treatment as indicated. SNRNP70 and HEXIM1 serve as positive controls for the U1 and 7SK snRNP complexes, respectively. **(P)** RT-qPCR analysis of the enrichment or depletion of the U1 or 7SK snRNAs from UV-crosslinked interfaces using either RNA antisense purification, or RNase H-mediated target RNA depletion (RNase H release) as implemented in RAP-MS or SHIFTR, respectively. Enrichment and depletion of target RNAs is normalized relative to GAPDH. Values are mean ± standard error of the mean. **(Q)** Deep proteome profile of 7SK SHIFTR experiments in Huh-7 (left) and A549 (right) cells using offline high pH reverse phase fractionation. Experiments shown correspond to data presented in Figures 1 E and G.

**Figure S2:**
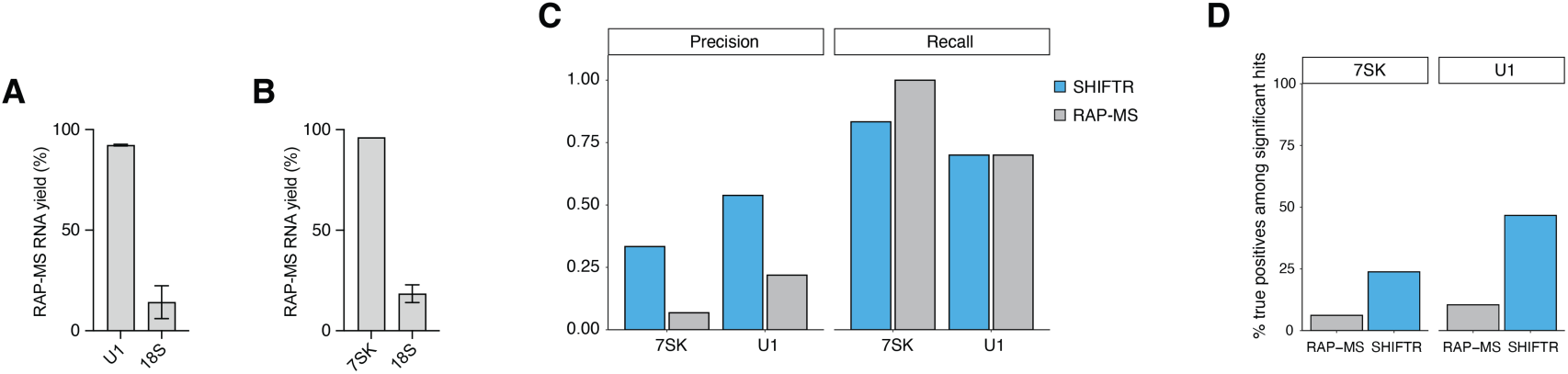
Side-by-side performance evaluation of SHIFTR and RAP-MS for identifying known U1 and 7SK components. **(A)** RT-qPCR analysis of RNA yield in RAP-MS experiments targeting the U1 snRNA. 18S rRNA serves as control. Yield is calculated relative to input. Values are mean ± standard error of the mean. **(B)** As in **(A)**, but for the 7SK snRNA. **(C)** Comparison of precision and recall statistics for uncovering known components of the U1 and 7SK snRNPs using either SHIFTR (blue) or RAP-MS (grey) to identify directly bound proteins. **(D)** Analysis of the percentage of true positive hits (known U1 or 7SK components) among all significantly enriched proteins in SHIFTR and RAP-MS experiments targeting the U1 and 7SK snRNPs.

**Figure S3:**
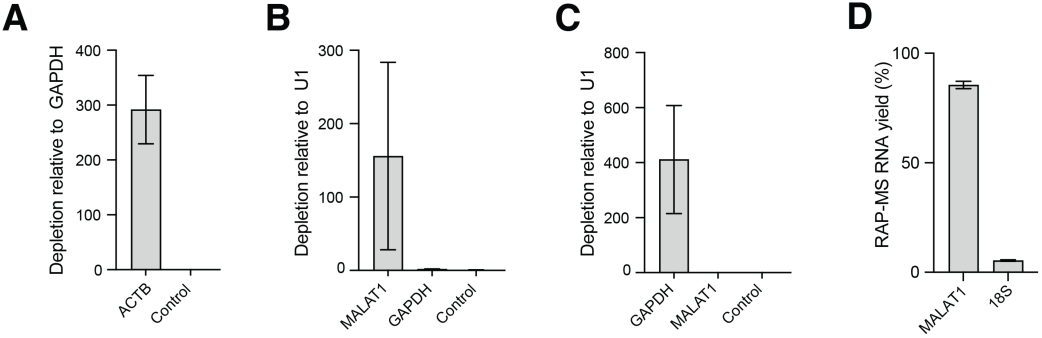
Targeting coding and non-coding RNAs with SHIFTR. **(A)** RT-qPCR analysis of RNase H-mediated depletion of the endogenous ACTB mRNA from UV-crosslinked interfaces using sequence-specific DNA probes. Depletion is normalized relative to GAPDH. Interface subjected to treatment without the addition of probes and enzyme serves as control. Values are mean ± standard error of the mean. **(B)** As in **(A)**, but targeting the MALAT1 lncRNA. Depletion is normalized relative to U1. Interfaces subjected to treatment with GAPDH-specific probes as well as interfaces subjected to treatment without the addition of probes and enzyme serve as controls. **(C)** As in **(B)**, but targeting the GAPDH mRNA. Interfaces subjected to treatment with MALAT1-specific probes as well as interfaces subjected to treatment without the addition of probes and enzyme serve as control. **(D)** RT-qPCR analysis of RNA yield in RAP-MS experiments targeting the MALAT1 lncRNA. Yield is calculated relative to input. 18S rRNA serves as control. Values are mean ± standard error of the mean.

**Figure S4:**
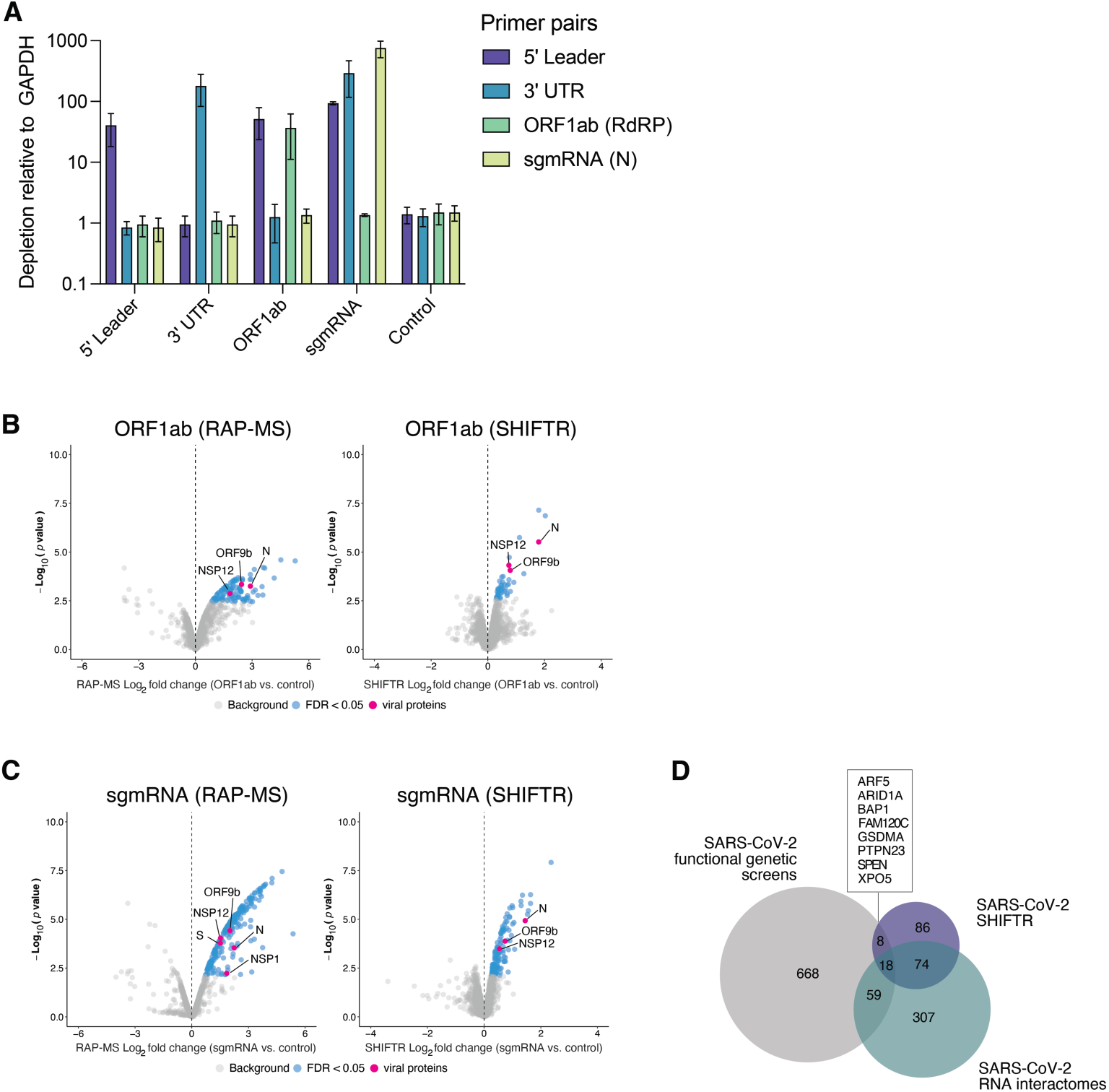
Interrogating different sequence regions in the authentic SARS-CoV-2 RNA genome with SHIFTR and RAP-MS. **(A)** RT-qPCR analysis of RNase H-mediated depletion of different regions in the SARS-CoV-2 RNA genome from UV-crosslinked interfaces using sequence-specific DNA probes. Interfaces isolated from infected human A549^ACE2^ cells (24 hpi) were used. Primer pairs targeting the indicated viral RNA regions (5ʹ leader, ORF1ab (RdRP), sgmRNA (N), 3ʹ UTR) were used and are indicated in different colors. Interfaces treated with sequence-specific DNA probes are indicated on the x-axis. Samples subjected to treatment without the addition of probes and enzyme serve as control. Depletion is normalized relative to GAPDH. Values are mean ± standard error of the mean. **(B)** Left: Quantitative analysis of proteins significantly enriched in purifications of SARS-CoV-2 RNA genomes using RAP-MS and targeting the ORF1ab region. Right: Quantitative analysis of proteins significantly enriched in SHIFTR experiments targeting the ORF1ab region in the SARS-CoV-2 RNA genome. Experiments were carried out in infected human cells at 24 hpi. Volcano plots display log_2_ fold changes and significance estimates for identified proteins. RAP-MS experiments utilize the unrelated RNA RMRP as the control. SHIFTR experiments utilize UV-crosslinked interfaces subjected to treatment without the addition of probes and enzyme as the control. Significantly enriched proteins are highlighted in blue, viral proteins are highlighted in pink. **(C)** As in **(B)**, but for SARS-CoV-2 sgmRNAs. RAP-MS experiments were first depleted for full length RNA genomes by capturing RNAs containing the ORF1ab region, prior to antisense-mediated capture of all sgmRNAs. **(D)** Venn diagram comparing hits identified in SARS-CoV-2 functional genetic screens (21 independent studies) (41) and SARS-CoV-2 RNA interactomes (5 independent studies) (41) with proteins significantly enriched in SHIFTR experiments targeting different SARS-CoV-2 RNA regions (5ʹ leader, ORF1ab, sgmRNAs, 3ʹ UTR).

## SUPPLEMENTARY TABLE LEGENDS

**Supplementary Table S1:** Quantitative mass spectrometry data related to Figure 1. SHIFTR optimization experiments targeting the endogenous U1 and 7SK snRNAs in two different cell types. Control data (no enzyme, no probe) is included in table as well.

**Supplementary Table S2:** Quantitative mass spectrometry data related to Supplementary Figure S1. SHIFTR optimization experiments recorded without offline high pH reverse phase fractionation. Control data (no enzyme, no probe) is included in table as well.

**Supplementary Table S3:** Quantitative mass spectrometry data related to Figure 2 and 3. Side-by-side comparison between SHIFTR and RAP-MS for different endogenous target RNAs.

**Supplementary Table S4:** Quantitative mass spectrometry data related to Figure 3. Multiplexed SHIFTR experiments targeting the endogenous ACTB mRNA and MALAT1 lncRNA are shown together with individual SHIFTR experiments targeting the GAPDH mRNA.

**Supplementary Table S5:** Quantitative mass spectrometry related to Figure 4. SHIFTR experiments targeting different sequence regions in the SARS-CoV-2 RNA genome (5ʹ leader, ORF1ab, sgmRNAs, 3ʹ UTR) are shown. GO enrichment analyses for each target RNA interactome (biological process (BP) and molecular function (MF)) is provided in separate worksheet.

**Supplementary Table S6:** Oligonucleotide sequences used in this study.

